# *E. coli* RecBCD Nuclease Domain Regulates Helicase Activity but not Single Stranded DNA Translocase Activity

**DOI:** 10.1101/2023.10.13.561901

**Authors:** Nicole Fazio, Kacey N. Mersch, Linxuan Hao, Timothy M. Lohman

**Author notes:** These authors contributed equally to this work (Nicole Fazio, Kacey N. Mersch).

## Abstract

Much is still unknown about the mechanisms by which helicases unwind duplex DNA. Whereas structure-based models describe DNA unwinding as a consequence of mechanically pulling the DNA duplex across a wedge domain in the helicase by the single stranded (ss)DNA translocase activity of the ATPase motors, biochemical data indicate that processive DNA unwinding by the *E. coli* RecBCD helicase can occur in the absence of ssDNA translocation of the canonical RecB and RecD motors. Here, we present evidence that dsDNA unwinding is not a simple consequence of ssDNA translocation by the RecBCD motors. Using stopped-flow fluorescence approaches, we show that a RecB nuclease domain deletion variant (RecB^ΔNuc^CD) unwinds dsDNA at significantly slower rates than RecBCD, while the rate of ssDNA translocation is unaffected. This effect is primarily due to the absence of the nuclease domain and not the absence of the nuclease activity, since a nuclease-dead mutant (RecB^D1080A^CD), which retains the nuclease domain, showed no significant change in rates of ssDNA translocation or dsDNA unwinding relative to RecBCD on short DNA substrates (≤ 60 base pairs). This indicates that ssDNA translocation is not rate-limiting for DNA unwinding. RecB^ΔNuc^CD also initiates unwinding much slower than RecBCD from a blunt-ended DNA, although it binds with higher affinity than RecBCD. RecB^ΔNuc^CD also unwinds DNA ∼two-fold slower than RecBCD on long DNA (∼20 kilo base pair) in single molecule optical tweezer experiments, although the rates for RecB^D1080A^CD unwinding are intermediate between RecBCD and RecB^ΔNuc^CD. Surprisingly, significant pauses occur even in the absence of *chi* (crossover hotspot instigator) sites. We hypothesize that the nuclease domain influences the rate of DNA base pair melting, rather than DNA translocation, possibly allosterically. Since the rate of DNA unwinding by RecBCD also slows after it recognizes a *chi* sequence, RecB^ΔNuc^CD may mimic a post-*chi* state of RecBCD.

## Introduction

*E. coli* RecBCD is a helicase/nuclease that functions in double stranded (ds)DNA break repair by binding to the DNA end, unwinding the DNA and loading RecA protein onto the resected 3’-single stranded (ss) DNA to initiate homologous recombination [1, 2]. RecBCD is composed of three subunits, RecB (134 kDa), RecC (129 kDa) and RecD (67 kDa) **(Fig. 1)**. RecB is a superfamily 1A helicase/ssDNA translocase that moves along ssDNA in the 3’ to 5’ direction, and RecD is a superfamily 1B helicase/ss DNA translocase that moves in the 5’ to 3’ direction [3–8]. During DNA unwinding, RecB and RecD translocate along the complementary ssDNA strands, such that both motors move in the same net direction [5, 6]. In addition to its canonical 3’ to 5’ translocase activity, the RecB motor also controls a secondary translocase activity involving the RecB arm domain, that may reflect a dsDNA translocase activity that may contribute to DNA base pair melting during DNA unwinding [7] [8–12].

**Figure 1.**
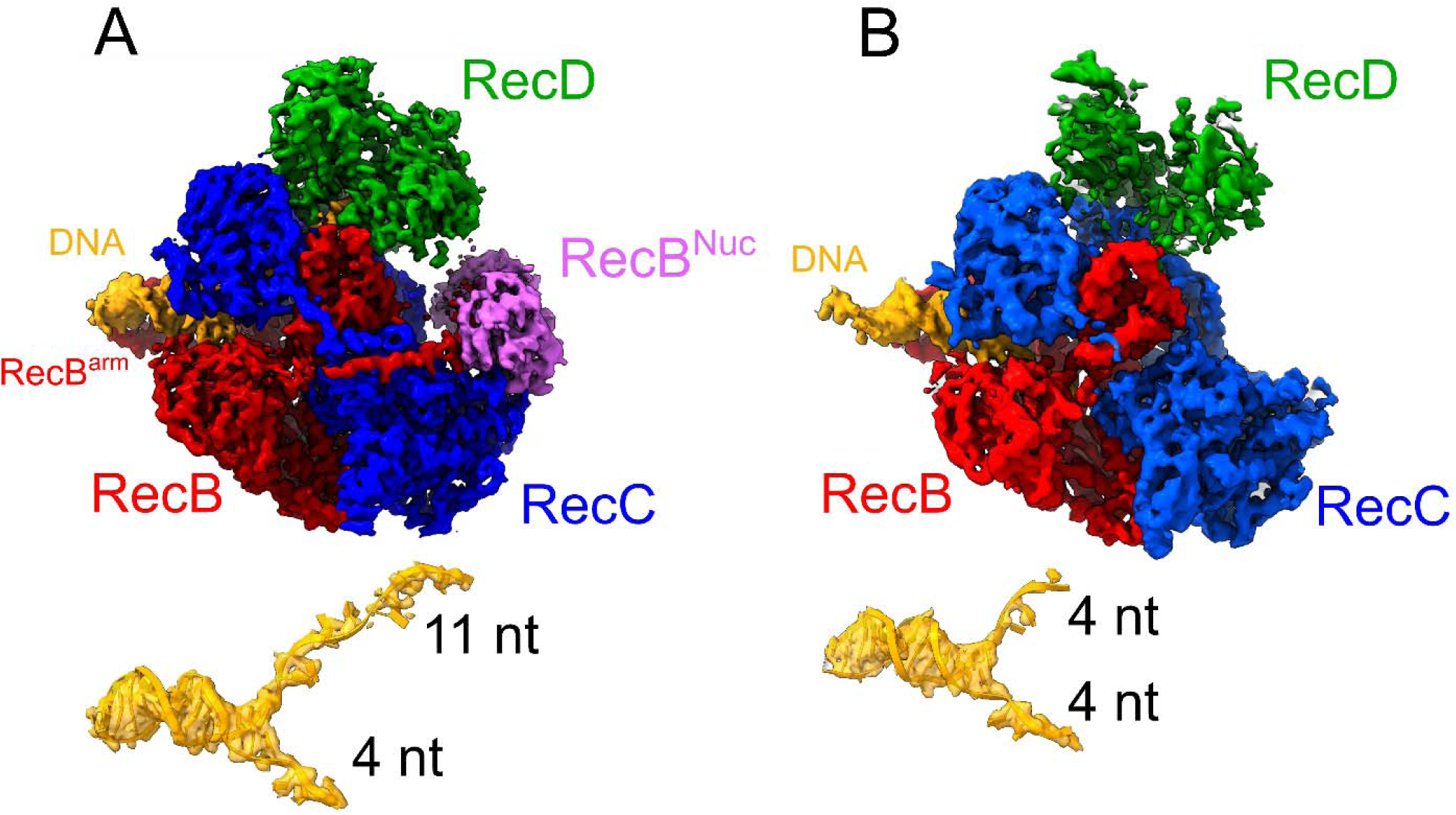
Two classes of cryo-EM structures of RecBCD bound to blunt-ended DNA. RecB motor domain (red), RecB nuclease domain (magenta), RecC (blue), RecD (green), and DNA (yellow). **(A)** RecBCD can melt up to 11 bp of DNA from a blunt end. Eleven bp are observed to be melted in one class of structures for which densities for the whole RecD subunit and RecB^Nuc^ are observed **(PDB: 7MR3)**[18]. **(B)** RecBCD melts only ∼4 bp from a blunt end in a second class of structures for which the RecB^Nuc^ and RecD 2B domain densities are not observed **(PDB: 7MR4)**[18].

The RecB subunit also contains a 30 kDa nuclease domain **(Fig. 1A)** that is responsible for DNA degradation during DNA unwinding [13–15]. The nuclease domain is attached to the RecB motor domain via a ∼60 amino acid linker, and in multiple RecBCD-DNA structures is observed in a docked position well removed from the duplex DNA binding site of RecBCD [16–18] **(Fig. 1A)**. RecC is structurally similar to RecB but does not have known motor activities [16, 19]. RecC interacts with both RecB and RecD and contains the site for recognition of the crossover hotspot instigator (*chi*) sequence (5’-GCTGGTGG-3’), which is an overrepresented regulatory sequence in the *E. coli* genome [20, 21]. Recognition of *chi* regulates the nuclease domain, affecting whether RecBCD initiates repair of the host DNA or continues to degrade foreign DNA, and also affects the rate of DNA unwinding [21, 22].

Initiation of repair of a dsDNA break occurs when RecBCD binds to a duplex DNA end and uses its binding free energy to melt as much as 4-11 bp [23] [16–18, 24] in the absence of ATP binding or hydrolysis **(Fig. 1)**. Upon ATP binding and hydrolysis, processive DNA unwinding begins. Prior to reaching a *chi*-site, the enzyme translocates faster along one strand in the 5’ to 3’ direction, suggesting that RecD is the faster motor during pre-*chi* unwinding [25]. This results in formation of a ssDNA loop in the other DNA strand ahead of the RecB motor [6], while the nuclease activity degrades both the 3’ and 5’ ssDNA ends [26]. Upon reaching and recognizing a *chi*-site, RecBCD pauses and undergoes a yet uncharacterized conformational change [27, 28]. Post *chi* recognition, RecBCD unwinds DNA at a ∼two-fold slower rate and the 3’ end of the DNA becomes sequestered within RecC so that the nuclease domain degrades only the 5’-ended DNA strand [25]. RecBCD then facilitates loading of RecA protein onto the resected 3’ single strand resulting in a RecA filament that initiates homologous recombination. The RecB nuclease domain has been proposed to interact directly with RecA protein to facilitate its loading onto ssDNA [29].

Since the proposed RecA binding site on the nuclease domain is occluded in the structures of RecBCD bound to DNA, it has been suggested that the nuclease domain is dynamic and may undock from the rest of RecBCD to facilitate RecA loading [29–32]. In fact, recent cryoEM structures of RecBCD bound to blunt-ended DNA show two classes of RecBCD-DNA structures [18]. In one class, density for the RecB nuclease domain is clearly present and up to 11 bp are melted from the blunt DNA end in an ATP-independent manner **(Fig. 1A)**. In a second class, RecBCD bound to the same blunt-ended DNA shows only ∼4 bp melted and no density for the nuclease domain is apparent **(Fig. 1B)**, suggesting that the nuclease domain has undocked [18].

Given that the rate of DNA unwinding and the nuclease activity are both affected post*-*chi, we hypothesized that the nuclease domain may regulate the rate of DNA unwinding. To examine this, we used a combination of ensemble stopped-flow fluorescence and single molecule optical tweezer experiments to compare the rates of ssDNA translocation and dsDNA unwinding of RecBCD and a variant in which the nuclease domain has been deleted (RecB^ΔNuc^CD) [8, 33, 34]. We show that removal of the nuclease domain reduces the rate of DNA unwinding by 25-45% compared to RecBCD. However, removal of the nuclease domain did not affect ssDNA translocation rates, implicating the nuclease domain as an allosteric regulator of DNA duplex melting. These observations support DNA unwinding models in which base pair melting is uncoupled from ssDNA translocation [35]. We suggest that RecB^ΔNuc^CD may mimic the post*-chi* state of RecBCD, and that undocking of the nuclease domain results in a slower DNA unwinding rate by slowing the rate of DNA base pair melting.

## Results

### DNA unwinding by stopped-flow fluorescence

We first examined the kinetics of dsDNA unwinding by RecBCD and RecB^ΔNuc^CD using an all-or-none single turnover fluorescence stopped-flow experiment as described [33, 34, 36] in buffer M at the indicated [NaCl] at 37°C. Duplex DNA substrates were labeled with a Cy3 and Cy5 FRET pair **(Fig. 2A, 3A)** and the Cy3 fluorescence was excited. Reactions were initiated by rapidly mixing pre-bound enzyme-DNA with ATP:Mg^2+^ and protein trap which binds any free enzyme ensuring single round DNA unwinding.

**Figure 2.**
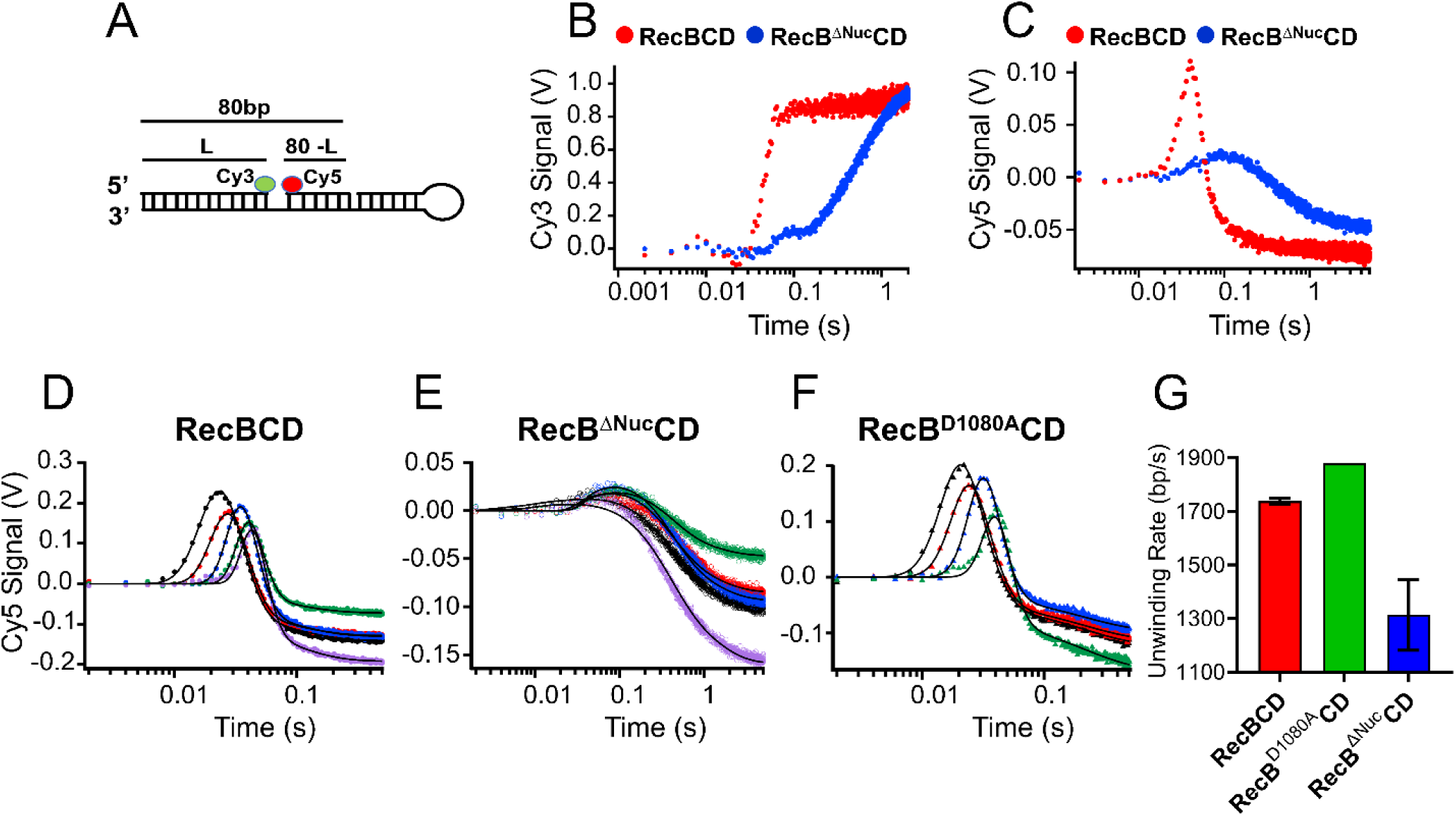
Representative stopped-flow time courses for RecBCD, RecB^Δ^^Nuc^CD, and RecB^D1080AC^D unwinding from a blunt DNA end (buffer M_30_ at 37°C, 5 mM ATP). **(A)** blunt-ended DNA substrate. **(B)** DNA unwinding traces monitoring the Cy3 fluorescence signal for RecBCD (red) and RecB^ΔNuc^CD (blue) (L = 50 bp). **(C)** DNA unwinding traces monitoring the Cy5 fluorescence signal for RecBCD (red) and RecB^ΔNuc^CD (blue) (L = 50 bp). Representative time courses for DNA unwinding of blunt ended DNA of L = 25 (black), 30 (red), 40 (blue), 50 (green), 60 (purple) bp by **(D)** RecBCD, **(E)** RecB^ΔNuc^CD, and **(F)** RecB^D1080A^CD. Global non-linear least square fits using Scheme 1 are shown as black solid lines. The best-fit parameters for the RecBCD data are: *k_U_* = 1306 s^−1^, *m* = 1.32 bp step^−1^, *k_NP_* = 12.3 s^−1^, *k_C_* = 151 s^−1^, h = 3.2 steps, x = 0.75, and *mk_U_* = 1726 bp s^−1^. The best-fit parameters for the RecB^ΔNuc^CD data are: *k* = 225 s^−1^, *m* = 5.30 bp step^−1^, *k_NP_* = 1.0 s^−1^, *k_C_* = 4 s^−1^, h = 1.3 steps, x = 0.62, and *mk_U_* = 1351 bp s^−1^. The best-fit parameters for the RecB^D1080A^CD data are: *k_U_* = 643 s^−1^, *m* = 2.92 bp step^−1^, *k_NP_* = 4.67 s^−1^, *k_C_* = 106 s^−1^, h = 1.6 steps, x = 0.59, and *mk_U_* = 1879 bp s^−1^. **(G)** Average macroscopic unwinding rates from blunt-ended DNA for RecBCD, RecB^ΔNuc^CD and RecB^D1080A^CD.

The all-or-none DNA unwinding time courses monitoring Cy3 fluorescence display a lag phase followed by an increase in Cy3 fluorescence due to a loss in FRET resulting from complete unwinding and DNA strand separation **(Fig. 2B, 3B)**. The lag phase reflects the time required for the enzyme to initiate, unwind the DNA and release the Cy3 labeled strand. The Cy5 fluorescence time courses **(Fig. 2C, 3C)** display a lag phase followed by a transient increase in fluorescence, which results from the transfer of a transient Cy3 fluorescence increase to the Cy5 fluorophore via FRET [33]. The Cy5 fluorescence then decreases as the Cy3 donor strand dissociates, resulting in a loss of FRET and low Cy5 fluorescence. The time-dependent changes in fluorescence signals were monitored for a series of DNA substrates that vary in DNA length, *L*, with *L* = 25, 30, 37, 40, 43, 48, 50, 53, and 60 bp (only traces for 25, 30, 40, 50, 60 bp are shown in the Figures for clarity).

### RecB^ΔNuc^CD initiates DNA unwinding from a blunt DNA end much slower than RecBCD

The stopped-flow time courses for unwinding of blunt ended DNA show significant qualitative differences for RecBCD and RecB^ΔNuc^CD (**Fig. 2B and 2C**; Buffer M_30_, 37°C, 5 mM ATP). When pre-bound to a blunt DNA end, RecBCD shows nearly complete unwinding by ∼80 ms after the addition of ATP, whereas RecB^ΔNuc^CD does not completely unwind the same length of DNA until ∼2 s after ATP addition **(Fig. 2B and C)**. The same final fluorescence is achieved for DNA unwinding by both RecBCD and RecB^ΔNuc^CD, suggesting that RecB^ΔNuc^CD unwinds the same amount of DNA, but more slowly. The Cy3 fluorescence time courses for both RecB^ΔNuc^CD and RecBCD show two phases **(Fig. 2B)**, but the slower phase is significantly larger for RecB^ΔNuc^CD. We interpret these two phases as arising from two conformational populations of RecBCD or RecB^ΔNuc^CD-DNA complexes ((*RD*)*_NP_* and (*RD*)*_P_*) in Scheme 1) prior to initiating unwinding, as discussed below. The populations of the two phases were calculated as described in Materials and Methods. Since the time courses monitoring the Cy5 fluorescence signal are more similar for RecBCD and RecB^ΔNuc^CD, we analyzed these time courses quantitatively to compare the kinetic parameters. The Cy3 and Cy5 signals have been shown to report on the same kinetic steps [33].

A series of time courses were collected for unwinding of blunt-ended DNA of varying lengths, *L*, by RecBCD or RecB^ΔNuc^CD. RecBCD shows the expected length-dependent lag phase **(Fig. 2D)**. In contrast, the lag phase for DNA unwinding by RecB^ΔNuc^CD shows less of a dependence on DNA length **(Fig. 2E)**. This suggests that a step preceding DNA unwinding is at least partially rate limiting for RecB^ΔNuc^CD [37].

We fit the Cy5 fluorescence time courses to the n-step kinetic model in Scheme 1 **(Eq. (2)**, and **(3))** using the non-linear least squares (NLLS) program MENOTR [38](see Materials and Methods). Scheme 1 has been shown to provide a good description of the kinetics of RecBCD unwinding dsDNA from a blunt end [33, 34, 36]. Figure 2D shows the best fit of Scheme 1 to a representative set of RecBCD-blunt end DNA unwinding time courses (**see Supplementary Table S2** for the kinetic parameters for individual data sets). The average best fit kinetic parameters for DNA unwinding from a blunt end are given in Table 1. The average macroscopic rate for RecBCD unwinding from a blunt DNA end is *mk_U_*= 1783 ± 11 bp s^−1^ (Buffer M_30_, 37°C, 5 mM ATP).

**Table 1.** Stopped-flow DNA unwinding kinetic parameters (Scheme 1) in Buffer M30, 37°C.

| Protein | DNA end type | $mk_U$ (bp s<sup>-1</sup>) | $k_U$ (s<sup>-1</sup>) | $m$ (bp step<sup>-1</sup>) | $k_{NP}$ (s<sup>-1</sup>) | $k_C$ (s<sup>-1</sup>) | $h$ (steps) | $x$ | slope |
| --- | --- | --- | --- | --- | --- | --- | --- | --- | --- |
| BCD | Blunt | 1738 ± 11 | 1150 ± 141 | 1.53 ± 0.19 | 9.7 ± 2.2 | 141 ± 10 | 2.7 ± 0.5 | 0.77 ± 0.02 | 0.6617 ± 0.0847 |
|  | T<sub>6</sub>T<sub>10</sub> | 1791 ± 36 | 1279 ± 202 | 1.43 ± 0.23 | 12.2 ± 0.9 | 113. ± 6 | 1.5 ± 0.3 | 0.74 ± 0.04 | 0.7148 ± 0.1200 |
| B<sup>Δnuc</sup>CD | Blunt | 1314 ± 132 | 460 ± 193 | 3.35 ± 1.77 | 1.0 ± 0.04 | 3.9 ± 0.4 | 1.3 ± 0.1 | 0.67 ± 0.05 | - |
|  | T<sub>6</sub>T<sub>10</sub> | 1238 ± 43 | 511 ± 148 | 2.57 ± 0.80 | 5.1 ± 0.4 | 29 ± 6 | 1.2 ± 0.2 | 0.46 ± 0.07 | - |
| B<sup>D1080A</sup>CD | Blunt | 1879 | 643 | 2.92 | 4.7 | 106 | 1.6 | 0.59 | 0.3423 |
|  | T<sub>6</sub>T<sub>10</sub> | 1701 | 897 | 1.90 | 8.1 | 139 | 1.5 | 0.61 | 0.5276 |

The Cy5 time courses for RecB^ΔNuc^CD unwinding from a blunt DNA end were also analyzed using Scheme 1. However, we used Eq. (3), employing the equivalency of *n* = *L/m* since the lag phase for RecB^ΔNuc^CD unwinding from a blunt DNA end showed only a slight dependence on DNA duplex length, such that the number of steps, *n*, could not be determined accurately for each DNA length. In Eq. (4), *L* is fixed for each length, and *m* is a global fitting parameter. Figure 2E shows the best fit of Scheme 1 to a representative data set for RecB^ΔNuc^CD-catalyzed unwinding from a blunt DNA end. The best fit kinetic parameters for the individual stopped-flow data sets are in **Supplementary Table S2, S3, S4, S7.** The average best fit parameters obtained from three sets of experiments are given in Table 1. The average macroscopic unwinding rate for RecB^ΔNuc^CD unwinding from a blunt DNA end is *mk*_U_ = 1314 ± 132 bp s^−1^.

These results suggest that RecB^ΔNuc^CD (1314 ± 132 bp/s) unwinds dsDNA from a blunt end significantly (∼25%) slower than RecBCD (1783 ± 11) **(Fig. 2E)**. In addition, the rate of initiation of dsDNA unwinding from a blunt end for RecB^ΔNuc^CD (*k*_C_ = 3.9 ± 0.4 s^−1^) is ∼40-fold slower than for RecBCD (*k*_C_ = 141 ± 10 s^−1^). The *k*_C_ step has been suggested to be associated with engagement of the RecD motor with the DNA [34], hence this step is significantly slower in the absence of the nuclease domain.

The NLLS analysis of the time courses also indicate that the rate of isomerization from non-productive, (*RD*)*_NP_*, to productive complexes, (*RD*)*p*, is nearly 10-fold slower for RecB^ΔNuc^CD (*k_NP_* = 1.0 s^−1^) compared to RecBCD (*k_NP_*= 9.7 s^−1^) **(Table 1)**. In addition, the fraction of complexes initially in the productive state, (*RD*) is much smaller for RecB^ΔNuc^CD than for RecBCD (**Table 1**)). Hence, the majority of RecB^ΔNuc^CD-DNA complexes are initially bound as non-productive complexes. These differences are not due to a lower affinity of RecB^ΔNuc^CD for a blunt DNA end. In fact, RecB^ΔNuc^CD binds with higher affinity to a blunt DNA end than does RecBCD [39].

### DNA unwinding from a pre-melted 3’-dT_6_/5’-dT_10_ DNA end

The slow rate of initiation of DNA unwinding for RecB^ΔNuc^CD from blunt-ended DNA limits the accuracy with which we can determine the macroscopic rate of DNA unwinding. We therefore also performed stopped-flow experiments with DNA substrates that possess a pre-melted (3’-dT_6_/5’-dT_10_) end (**Fig. 3A**). RecBCD binds with high affinity to such a “pre-melted” initiation site, and the 3’ and 5’ DNA ends are sufficiently long to engage the RecB and RecD motors, respectively [18, 34, 40]. With these DNA substrates, the unwinding time courses for RecB^ΔNuc^CD (and RecBCD) show clear lag phases that increase with DNA length indicating that the slow rate of initiation for RecB^ΔNuc^CD is eliminated (**Fig. 3E**). The Cy3 DNA unwinding time course for RecB^ΔNuc^CD still shows a slow second phase beginning ∼0.1-0.15 s (**Fig. 3B and C**), but this slow phase has a smaller amplitude than for initiation from a blunt DNA end (**Fig. 2B**).

**Figure 3.**
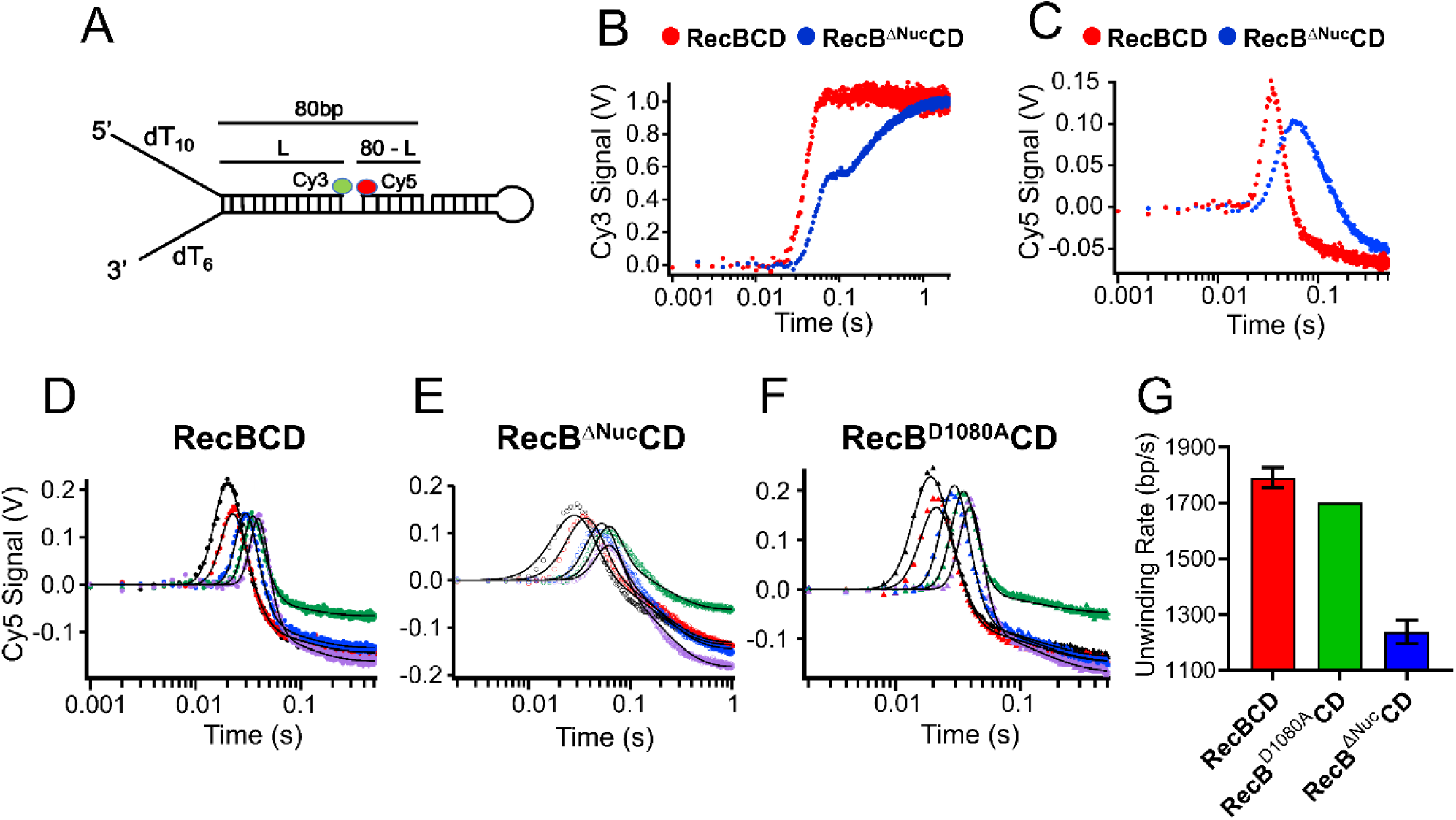
Representative stopped-flow time courses for RecBCD, RecB^ΔNuc^CD, and RecB^D1080AC^D unwinding from a pre-melted 3’-dT_6_/5’-dT_10_ DNA end (buffer M_30_ at 37°C, 5 mM ATP). **(A)** Pre-melted 3’-dT_6_/5’-dT_10_ DNA substrate. DNA unwinding traces monitoring the Cy3 fluorescence signal for RecBCD (red) and RecB^ΔNuc^CD (blue) (L = 50 bp). **(C)** DNA unwinding traces monitoring the Cy5 fluorescence signal for RecBCD (red) and RecB^ΔNuc^CD (blue) (L = 50 bp). Representative time courses for DNA unwinding of blunt ended DNA of L = 25 (black), 30 (red), 40 (blue), 50 (green), 60 (purple) bp by **(D)** RecBCD, **(E)** RecB^ΔNuc^CD, and **(F)** RecB^D1080A^CD. Global non-linear least square fits using Scheme 1 are shown as black solid lines. The best-fit parameters for the RecBCD data are: *k_U_* = 1105 s^−1^, *m* = 1.64 bp step^−1^, *k_NP_* = 11.3 s^−1^, *k_C_* = 108 s^−1^, h = 1.2 steps, x = 0.73, and *mk_U_* = 1813 bp s^−1^. The best-fit parameters for the RecB^ΔNuc^CD are: *k* = 669 s^−1^, *m* = 1.83 bp step^−1^, *k_NP_* = 4.8 s^−1^, *k_C_* = 29 s^−1^, h = 1.1 steps, x = 0.44, and *mk_U_* = 1226 bp s^−1^. The best-fit parameters for the RecB^D1080A^CD are: *k_U_* = 897 s^−1^, *m* = 1.90 bp step^−1^, *k_NP_* = 8.10 s^−1^, *k_C_* = 139 s^−1^, h = 1.5 steps, x = 0.61, and *mk_U_* = 1701 bp s^−1^. **(G)** Average macroscopic unwinding rates from blunt-ended DNA for RecBCD, RecB^ΔNuc^CD and RecB^D1080A^CD.

For consistency, we also analyzed the Cy5 time courses using Scheme 1 (**Eq. 3**). Figure 3D shows the best fit of Scheme 1 to a representative data set for RecBCD-catalyzed unwinding from a (3’-dT_6_/5’-dT_10_) end. The best fit kinetic parameters are reported in the Figure legend and the average best fit parameters from three data sets are given in **Table 1** (Rate constants from each individual data set are given in **Supplementary Table S2**. The average macroscopic rate of DNA unwinding for RecBCD is *mk_U_*= 1791 ± 36 bp/s.

Figure 3E shows the best fit of Scheme 1 to a representative data set for RecB^ΔNuc^CD-catalyzed unwinding from a (3’-dT_6_/5’-dT_10_) end. The average best fit parameters from three data sets are given in Table 1. The average macroscopic rate of DNA unwinding for RecBCD is *mk*_U_ = 1238 ± 43 bp/s. These results support the conclusion that RecB^ΔNuc^CD unwinds dsDNA ∼30% slower than does RecBCD.

The rate of initiation of unwinding for RecB^ΔNuc^CD is also slower than for RecBCD, but the difference in *k*_C_ is not as great as for unwinding from a blunt DNA end. The values of *k_NP_*and *k_C_* change only slightly for RecBCD unwinding from a pre-melted 3’-dT_6_/5’-dT_10_ DNA end vs. a blunt DNA end **(Table 1)**. However, the values of *k_NP_* and *k_C_* increase significantly (∼five-fold and ten-fold, respectively) for RecB^ΔNuc^CD unwinding from a pre-melted DNA end, compared to a blunt DNA end. Additionally, a greater fraction of productive RecB^ΔNuc^CD-DNA complexes are observed for unwinding from a pre-melted end. Hence, deletion of the nuclease domain significantly affects initiation from different DNA end types, but differences in initiation rates alone do not account for the slower DNA unwinding rates for RecB^ΔNuc^CD.

### RecB^ΔNuc^CD unwinds DNA slower than RecBCD at all salt concentrations

The DNA unwinding experiments described above were performed in buffer containing 30 mM NaCl. However, for technical reasons, the ssDNA translocation experiments described below were performed at 275 mM and 500 mM NaCl. Hence, we also examined DNA unwinding at these higher [NaCl] using DNA substrates with a 3’-dT_6_/5’-dT_10_ end **(Tables 2 and 3, Supplementary Figures S1 and S2 and Tables S3 and S4)**. Four sets of time courses were obtained for both RecBCD and RecB^ΔNuc^CD at 275 mM NaCl (buffer M_275_) and two sets of time courses were obtained at 500 mM NaCl (buffer M_500_) (37°C). These time courses were fit to Scheme 1 and the average kinetic parameters are given in Tables 2 and 3. In buffer M_275_, RecB^ΔNuc^CD unwinds with an average macroscopic rate of 1249 ± 36 bp/s that is 27% slower than RecBCD (1702 ± 21 bp/s). In buffer M_500_, RecB^ΔNuc^CD (858 ± 42 bp/s) unwinds ∼44% slower compared to RecBCD (1532 ± 17 bp/s). Hence, under all solution conditions tested, deletion of the nuclease domain results in a slower rate of DNA unwinding.

**Table 2.** Stopped-flow DNA unwinding kinetic parameters (Scheme 1) in Buffer M275, 37°C.

| Protein | DNA end type | $mk_U$ (bp s<sup>-1</sup>) | $k_U$ (s<sup>-1</sup>) | $m$ (bp step<sup>-1</sup>) | $k_{NP}$ (s<sup>-1</sup>) | $k_C$ (s<sup>-1</sup>) | $h$ (steps) | $x$ | slope |
| --- | --- | --- | --- | --- | --- | --- | --- | --- | --- |
| BCD | T<sub>6</sub>T<sub>10</sub> | 1702 ± 21 | 1148 ± 117 | 1.49 ± 0.15 | 10.3 ± 1.7 | 141 ± 30 | 1.3 ± 0.4 | 0.80 ± 0.06 | 0.6740 ± 0.0630 |
| B<sup>Δnuc</sup>CD |  | 1249 ± 36 | 734 ± 181 | 1.78 ± 0.46 | 5.2 ± 0.3 | 25 ± 5 | 1.1 ± 0.1 | 0.44 ± 0.04 | 0.5858 ± 0.1347 |
| B<sup>D1080A</sup>CD |  | 1698 ± 40 | 1043 ± 67 | 1.63 ± 0.14 | 8.4 ± 0.5 | 126 ± 11 | 1.1 ± 0.1 | 0.64 ± 0.03 | 0.6147 ± 0.0200 |

**Table 3.** Stopped-flow DNA unwinding kinetic parameters (Scheme 1) in Buffer M500, 37°C.

| Protein | DNA end type | $mk_U$ (bp s<sup>-1</sup>) | $k_U$ (s<sup>-1</sup>) | $m$ (bp step<sup>-1</sup>) | $k_{NP}$ (s<sup>-1</sup>) | $k_C$ (s<sup>-1</sup>) | $h$ (steps) | $x$ | slope |
| --- | --- | --- | --- | --- | --- | --- | --- | --- | --- |
| BCD | T<sub>6</sub>T<sub>10</sub> | 1532 ± 17 | 1368 ± 109 | 1.12 ± 0.10 | 10.1 ± 0.3 | 187 ± 12 | 2.4 ± 0.1 | 0.61 ± 0.01 | 0.8936 ± 0.0810 |
| B<sup>Δnuc</sup>CD |  | 858 ± 42 | 440 ± 405 | 3.30 ± 0.29 | 5.7 ± 4.0 | 19 ± 11 | 1.1 ± 0.1 | 0.39 ± 0.37 | 0.5021 ± 0.4473 |
| B<sup>D1080A</sup>CD |  | 1600 ± 1 | 1224 ± 123 | 1.31 ± 0.13 | 7.1 ± 0.3 | 215 ± 58 | 3.2 ± 1.4 | 0.49 ± 0.03 | 0.7653 ± 0.0774 |

During DNA unwinding by RecBCD, the unwound ssDNA is degraded by the nuclease activity of RecBCD unless a single stranded DNA binding protein, such as *E. coli* SSB, is present to bind and protect the unwound ssDNA[41, 42]. Since deletion of the nuclease domain results in the loss of nuclease activity, we considered whether the slower DNA unwinding rate of RecB^ΔNuc^CD might be related to the loss of nuclease activity. We therefore examined dsDNA unwinding by RecB^D1080A^CD, containing a single mutation (D1080A) in the nuclease active site that eliminates nuclease activity [43]. We find that the DNA unwinding rates for RecB^D1080A^CD initiating from either a blunt end **(Fig. 2F)** or a 3’(dT)_6_/5’(dT)_10_ end **(Fig. 3F)** do not differ significantly from those of RecBCD in buffer M_30_ at 37°C **(Figure 2G, Figure 3G, Table 1)**.

This is also the case for DNA unwinding at 275 mM NaCl and 500 mM NaCl **(Tables 2 and 3, Supplementary Fig. S1 and S2)**. Hence, the slower rate observed for RecB^ΔNuc^CD unwinding of the short DNA substrates in the stopped-flow experiments is not due to the loss of nuclease activity.

### Single molecule DNA unwinding rates measured by optical trapping

We also examined unwinding of single DNA molecules using a combined optical tweezer/fluorescence instrument (LUMICKS C-Trap G2) to examine the DNA unwinding rates for RecBCD, RecB^ΔNuc^CD and RecB^D1080A^CD. This approach allows a direct measurement of the macroscopic DNA unwinding rate but does not allow determination of the other kinetic parameters that can be assessed in the ensemble stopped-flow approach. The experimental setup (see **Materials and Methods**) is essentially as described [44] and is shown schematically in **Figure 4A** and **Supplemental Figure S3**. These experiments utilize a 19,435 bp dsDNA created from a 20,452 bp dsDNA that contained both a 3’- and 5’-5x-biotin tag on one strand of the dsDNA construct. A restriction endonuclease, SmaI, was used to cleave the 5’-5x-biotin tag to create a blunt DNA end to which the helicase can initiate DNA unwinding. The DNA end with the 3’-5x-biotin tag is bound to a streptavidin-coated polystyrene bead. We note that this DNA does not contain any *chi* sequences. The dsDNA was extended under flow in Buffer M_30_ containing Sytox Orange (SxO), a fluorophore that remains relatively dark in solution but undergoes a significant fluorescence increase upon DNA binding [45, 46]. Upon movement of the DNA to a channel containing RecBCD (100 nM) and ATP (2 mM), DNA unwinding was initiated and monitored by the release of the SxO dye. The rate of DNA unwinding is measured as a shortening of the duplex DNA through the action of the helicase **(Fig. 4B)**. We observed two classes of DNA unwinding trajectories; those that exhibit uninterrupted unwinding events **(Fig. 4B)**, as reported previously [25, 28, 44, 47], but also trajectories that exhibit significant pauses **(Figure 5)**.

**Figure 4.**
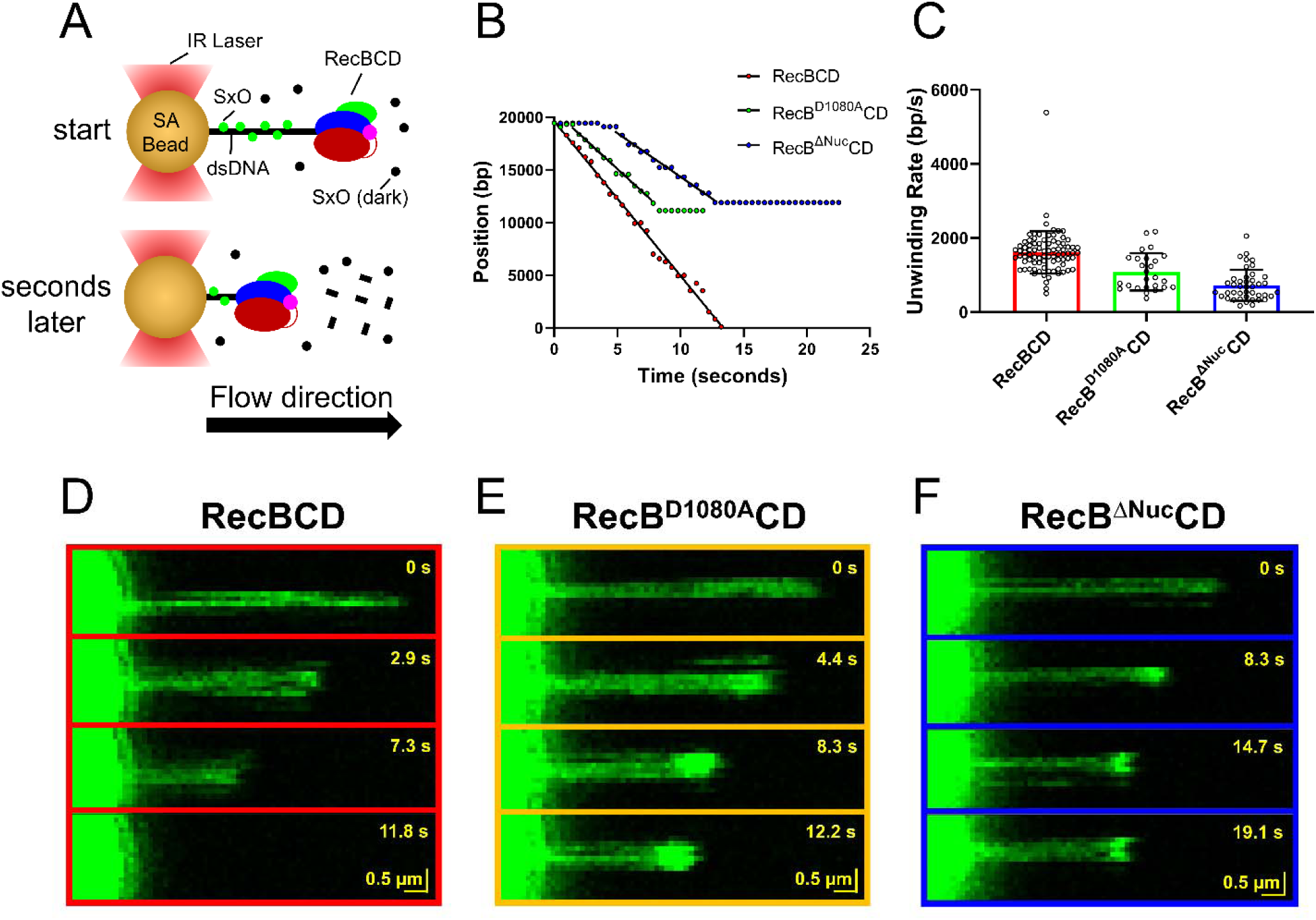
Single molecule DNA unwinding by RecBCD, RecB^ΔNuc^CD and RecB^D1080A^CD. **(A)** In a LUMICKS C-Trap, RecBCD and RecBCD variants are bound to a blunt-ended 19,435 bp DNA with the other end attached to a 4.34 μm streptavidin (SA) coated bead via a biotin-SA linkage in the presence of Sytox orange (Sxo) fluorophore that binds duplex DNA. In the presence of ATP, RecBCD starts to unwind the DNA, releasing the Sxo fluorophore. In the case of RecBCD, the newly formed single-stranded DNA is also degraded by its nuclease domain (magenta). **(B)** Representative plots showing the time dependence of the shortening of the duplex DNA lengths to obtain unwinding rates for RecBCD (1676 bp/s), RecB^D1080A^CD (1099 bp/s), and RecB^ΔNuc^CD (829 bp/s). These data show trajectories that do not contain significant pauses. **(C)** Rates of double-stranded DNA unwinding estimated only for trajectories that do not show pauses: RecBCD: 1612 ± 572 bp/s (n = 84 molecules), RecB^D1080A^CD: 1084 ± 503 bp/s (n = 26), and RecB^ΔNuc^CD: 723 ± 420 bp/s (n = 42) (mean ± SD (n)). Student’s T-test reveal that both RecB^D1080A^CD & RecB^ΔNuc^CD unwind DNA significantly slower than WT RecBCD (p-value <0.0001) and that RecB^ΔNuc^CD unwinds DNA significantly slower than RecB^D1080A^CD (p-value = 0.0022). **(D), (E), and (F)** Time-lapse of enzyme mediated shortening of DNA by **(D)** - RecBCD, **(E)** - RecB^D1080A^CD, and **(F)** - RecB^ΔNuc^CD.

**Figure 5.**
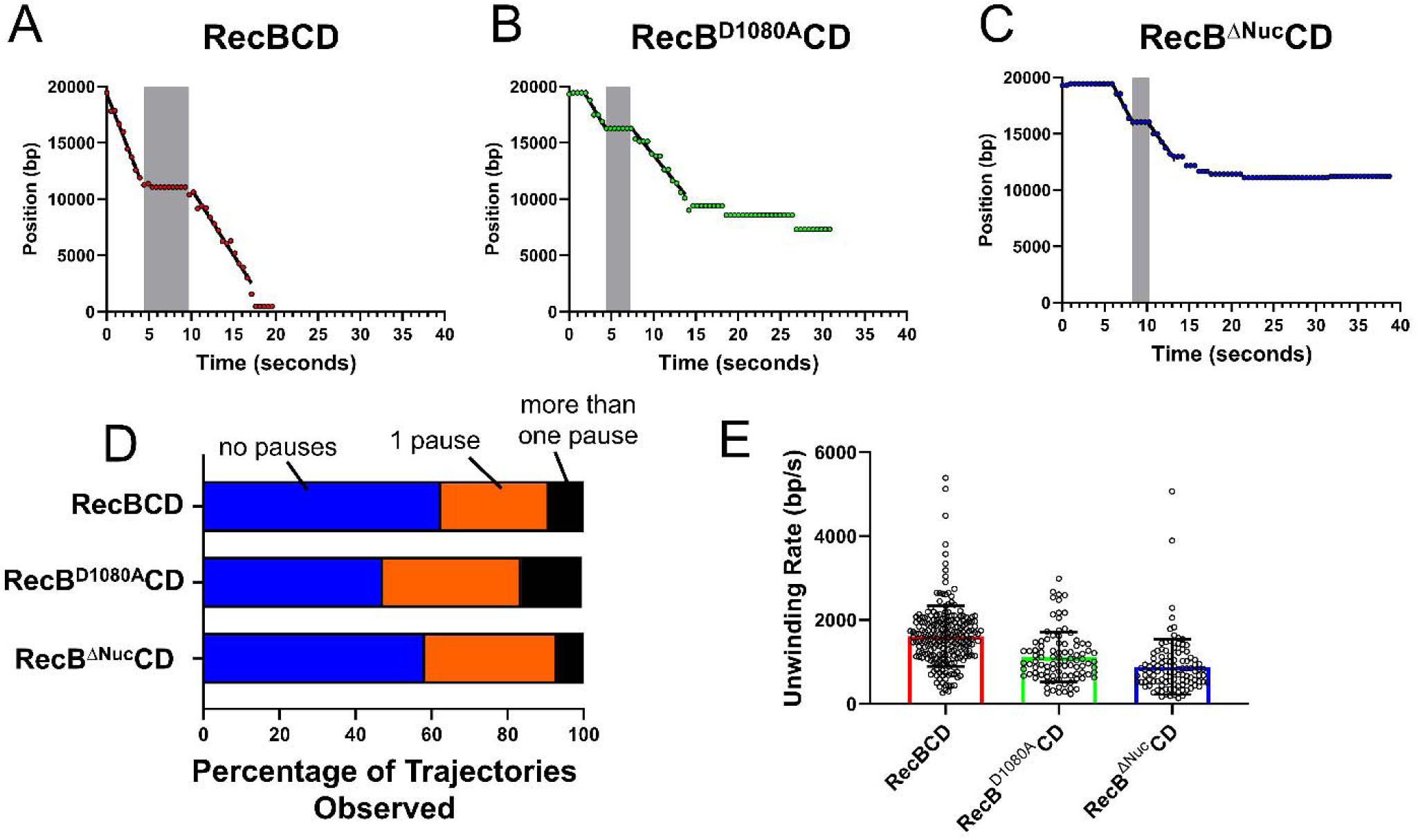
RecBCD and RecBCD variants show *chi-*independent pausing during DNA unwinding. Single DNA unwinding experiments performed on a 19,435 bp DNA with a LUMICKS C-Trap **(A), (B) and (C)** Examples of duplex DNA shortening trajectories displaying a single pause (shaded in gray) in DNA unwinding for **(A)** – RecBCD, **(B)** - RecB^D1080A^CD, and **(C)** - RecB^ΔNuc^CD. **(D)** The percentage of DNA unwinding trajectories for RecBCD (n = 144), RecB^D1080A^CD (n = 55), and RecB^ΔNuc^CD (n = 72) containing no pauses (blue), one pause (orange), and more than one pause (black). **(E)** DNA unwinding rates estimated from all data sets with and without pauses: RecBCD 1615 ± 721 bp/s (e= 218 unwinding events, n = 144 molecules), RecB^D1080A^CD 1117 ± 596 bp/s (e= 95, n = 55), and RecB^ΔNuc^CD 883 ± 658 bp/s (e= 109, n = 72 molecules) (mean ± SD bp/s.

Approximately 50-60% of the DNA unwinding trajectories showed no apparent pauses. Representative time-lapse recordings of RecBCD, RecB^ΔNuc^CD and RecB^D1080A^CD catalyzed DNA unwinding without pauses are shown in **Figure 4D, E** and **F**. DNA unwinding rates were determined from a linear least squares fit to the shortening trajectory before the cessation of DNA unwinding. Analysis of multiple single molecule trajectories showed average RecBCD rates of 1612 ± 572 bp/s vs. average RecB^ΔNuc^CD rates of 723 ± 420 bp/s (mean ± SD) **(Fig. 4C, Supplementary Table S5)**. Hence, the average rate of DNA unwinding for RecB^ΔNuc^CD is ∼45% slower than for RecBCD. The slower rate of DNA unwinding by RecB^ΔNuc^CD compared to RecBCD is consistent with the results from the ensemble stopped-flow studies. Interestingly, the average DNA unwinding rates for RecB^D1080A^CD (1084 ± 503 bp/s) were faster than for RecB^ΔNuc^CD, but slower than RecBCD **(Fig. 4C, Supplementary Table S5)**.

In contrast to the single molecule movies for RecBCD **(Fig. 4D)**, the movies for RecB^ΔNuc^CD **(Fig. 4E)** and RecB^D1080A^CD **(Fig. 4F)** show high intensity fluorescence spots near the end of the DNA that is being unwound. As suggested previously [28], these bright spots likely represent the formation of some secondary structure (base pairing) by the unwound single stranded DNA since both RecB^ΔNuc^CD and RecB^D1080A^CD lack nuclease activity of RecBCD that degrades the unwound single strands during DNA unwinding. The high intensity fluorescence spots are only present during DNA unwinding by RecB^ΔNuc^CD and RecB^D1080A^CD (**Supplemental Fig. S4**), in the absence of a single stranded DNA binding protein (see below).

We also observed a second class of DNA unwinding trajectories for all three RecBCD variants, that exhibit significant pausing during DNA unwinding (**Fig. 5A-C**). Approximately 30-40% of the trajectories exhibit a single pause, but trajectories showing multiple pauses also occur (∼5-10%) **(Fig. 5D)**. In trajectories with pauses, we estimated DNA unwinding rates by linear least squares fitting of the DNA unwinding trajectories between the pauses. These rates show the same trends in DNA unwinding rates as determined from the trajectories without pauses, with RecBCD rates (1615 ± 721 bp/s) nearly twice as fast as RecB^ΔNuc^CD (883 ± 658 bp/s)), and with RecB^D1080A^CD showing intermediate DNA unwinding rates (1117 ± 596 bp/s) **(Fig. 5E, Supplementary Table S5, and Supplemental Fig. S5)**. The duration of the pauses are between 5.8 and 7.7 seconds **(Supplemental Figure S6 A-C)**. We note that the duration of time each enzyme spends unwinding DNA (between pauses) (4.8-5.6 seconds), is similar to the duration time of the pauses **(Supplemental Fig. S6 D-F)**. These pauses are not *chi-*dependent since the DNA used in the LUMICKS experiments does not contain any *chi* sites. Furthermore, the pauses occur randomly and are not associated with any *chi* or *chi*-like DNA sequences **(Supplemental Fig. S7 A-C)**.

The DNA unwinding rates before and after these *chi*-independent pauses changed randomly **(Supplemental Fig. S8 A-C**. Previous experiments describe similar stochastic changes in unwinding rates after pauses in RecBCD unwinding induced by abrupt depletion of ATP or Mg^2+^[47]. This contrasts with the observation that the rate of DNA unwinding by RecBCD is slower after it encounters a *chi*-site [1, 28].

Although there are significant differences in the macroscopic unwinding rates of RecB^ΔNuc^CD and RecB^D1080A^CD compared to RecBCD **(Fig. 5E)**, each enzyme is highly processive and capable of unwinding thousands of base pairs **(Supplemental Fig. S9 and Supplementary Table S6)**. By determining when 50% of the RecBCD, RecB^ΔNuc^CD, and RecB^D1080A^CD enzymes have dissociated from the DNA, we can estimate a minimum average number of bp unwound by each enzyme: 21,165 bp for RecBCD, 14,192 bp for RecB^D1080A^CD, and 12,823 bp for RecB^ΔNuc^CD **(Supplementary Fig. S9 and Table S6)**.

### Effects of SSB protein on DNA unwinding

In the RecBCD DNA unwinding experiments shown in Figure 4, the nuclease activity of RecBCD degrades the single stranded DNA during DNA unwinding. However, this does not occur during DNA unwinding by RecB^ΔNuc^CD or RecB^D1080A^CD. Therefore, we also performed single molecule DNA unwinding experiments in the presence of *E. coli* SSB protein (**Supplementary Figure S10A**) that will bind with high affinity to the newly formed ssDNA inhibiting the nuclease activity of RecBCD [48]. The relative rates of DNA unwinding of the RecBCD variants in the presence of SSB protein show the same trends as in the absence of SSB protein **(Supplemental Figure S10B and Supplementary Table S5**) with RecBCD (2035 ± 278 bp/s) showing a ∼two-fold faster rate of unwinding than RecB^ΔNuc^CD (1033 ± 711 bp/s) with an intermediate rate for RecB^D1080A^CD (1270 ± 711 bp/s). It is noteworthy that rates of DNA unwinding are enhanced significantly in the presence of SSB protein **(Supplementary Table S5).** The experiments in the presence of SSB also show two classes of DNA unwinding trajectories, some with no pauses and others with one or more pauses (**Supplemental Figure S10C**). Notably, RecBCD showed fewer pauses in the presence of SSB protein (37% vs 23% of the trajectories). The changes in the rates of unwinding after these *chi-*independent pauses in the presence of *E. coli* SSB is stochastic (**Supplemental Figure S11**).

The durations of the pauses in the presence of SSB protein are in the range of 2.3-3.8 seconds, slightly smaller compared to those in the absence of SSB **(Supplemental Figure S12)**. The pausing durations are similar to the duration of the unwinding events, which range from 3.7-4.6 seconds. The data suggest that processivity is slightly enhanced in the presence of SSB protein (**Supplementary Figure S13; Supplementary Table S6**). Lastly, the inclusion of SSB protein nearly eliminates the bright fluorescence spot at the DNA end observed during unwinding by RecB^D1080A^CD and RecB^ΔNuc^CD **(Supplemental Figure S10D, S10E, and S4)**. Altogether, these data suggest that SSB slightly enhances the rate and processivity of DNA unwinding of RecBCD, RecB^D1080A^CD, and RecB^ΔNuc^CD and inhibits the formation of secondary structure in the unwound ssDNA.

### Single stranded DNA translocation rates

Processive DNA unwinding by a helicase requires that the helicase (1) melt or separate some number of DNA base pairs and (2) translocate directionally along the DNA. We have shown that deletion of the nuclease domain significantly slows the rate of dsDNA unwinding. To test whether this is due to a slower translocation rate, we compared the rates of ssDNA translocation for RecBCD, RecB^ΔNuc^CD and RecB^D1080A^CD using a stopped-flow fluorescence assay as described [7, 8]. The DNA substrates **(Fig. 6A and 7A)** used to measure ssDNA translocation have a 3’(dT)_6_/5’(dT)_10_ high affinity binding site at one end of a 24 bp duplex, followed by twin ssDNA (dT)_L_ tails of identical length, *L*, after the DNA duplex. To examine 3’ to 5’ ssDNA translocation, a Cy3 label is positioned at the 5’-terminated tail **(Fig. 6A)**. To examine 5’ to 3’ ssDNA translocation, a Cy3 label is positioned at the 3’-terminated tail **(Fig. 7A)**. The enzyme first binds at the high affinity binding site, then upon addition of ATP, it unwinds the 24 bp duplex and then continues to translocate along both ssDNA (dT)*_L_* tails until it reaches the Cy3 fluorophore resulting in enhancement of the Cy3 fluorescence [7, 8, 49, 50]. The Cy3 fluorescence enhancement is followed by a decrease in Cy3 fluorescence as the protein dissociates from the DNA end. By performing such experiments on multiple DNA substrates with varying lengths, *L*, one can obtain estimates of the ssDNA translocation rates as described [7, 8].

**Figure 6.**
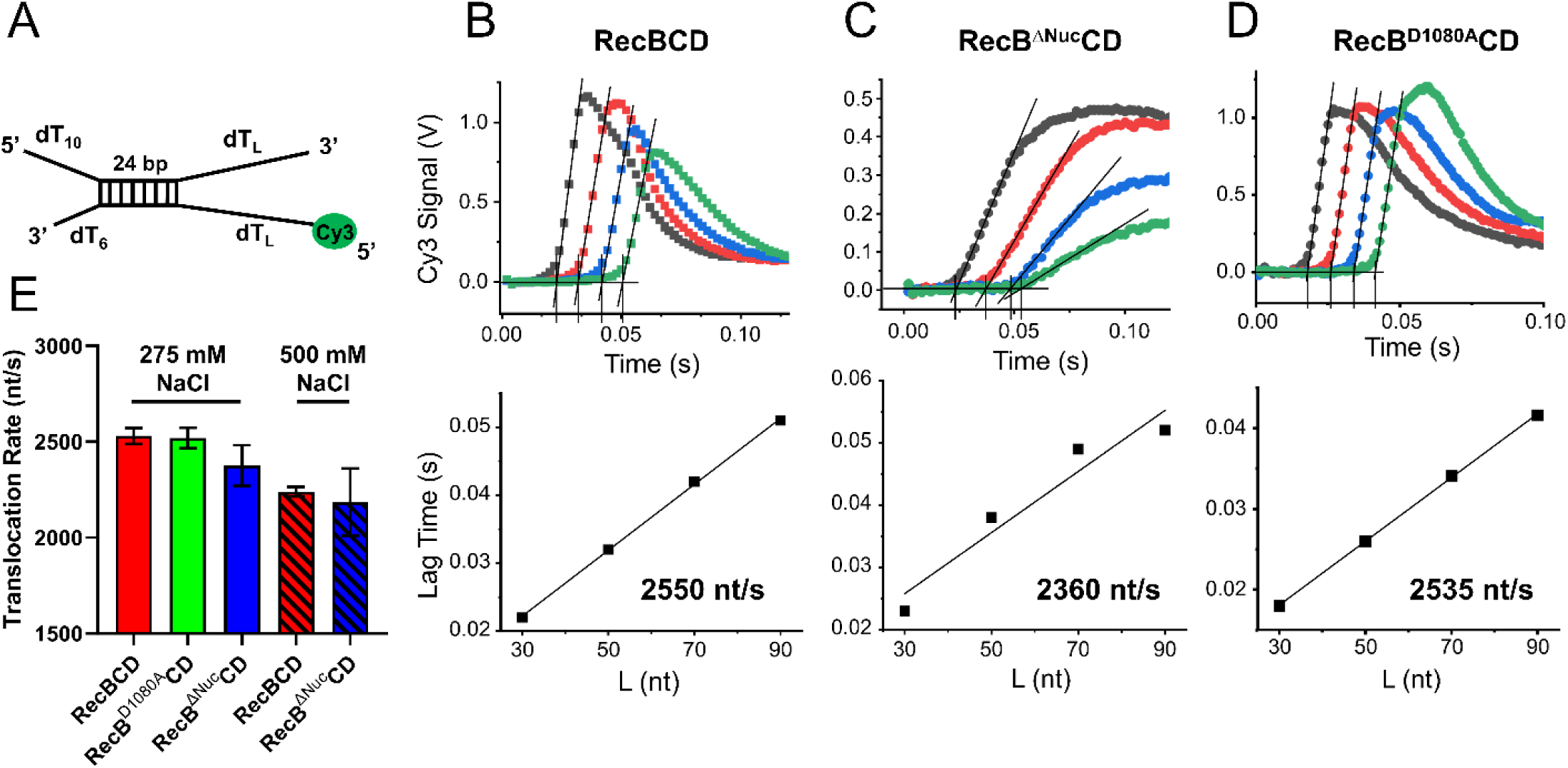
Stopped-flow time courses monitoring 3’ to 5’ ssDNA translocation of RecBCD, RecB^D1080A^CD, and RecB^ΔNuc^CD (M_275_ at 37°C, 5 mM ATP). **(A)** Pre-melted DNA substrate used to measure 3’ to 5’ ssDNA translocation. Representative DNA translocation time courses monitoring the enhancement of Cy3 fluorescence occurring when the helicase reaches the 5’ end of the ssDNA for **(B)** - RecBCD, **(C)** - RecB^D1080A^CD, and **(D)** - RecB^ΔNuc^CD. Time courses were obtained as a function of the 5’-ssDNA tail length (*L* = 30 nt (black); 50 nt (red); 70 nt (blue); 90 nt (green)). Lag times are plotted as a function of (L(nt)) below each data set and the macroscopic ssDNA translocation rate is estimated from the inverse of the slope of the best-fit line. **(E)** Average 3’ to 5’ ssDNA translocation rates for RecBCD, RecB^ΔNuc^CD, and RecB^D1080A^CD in buffer M_275_ (solid) and buffer M_500_ (striped) at 37°C and 5 mM ATP.

**Figure 7.**
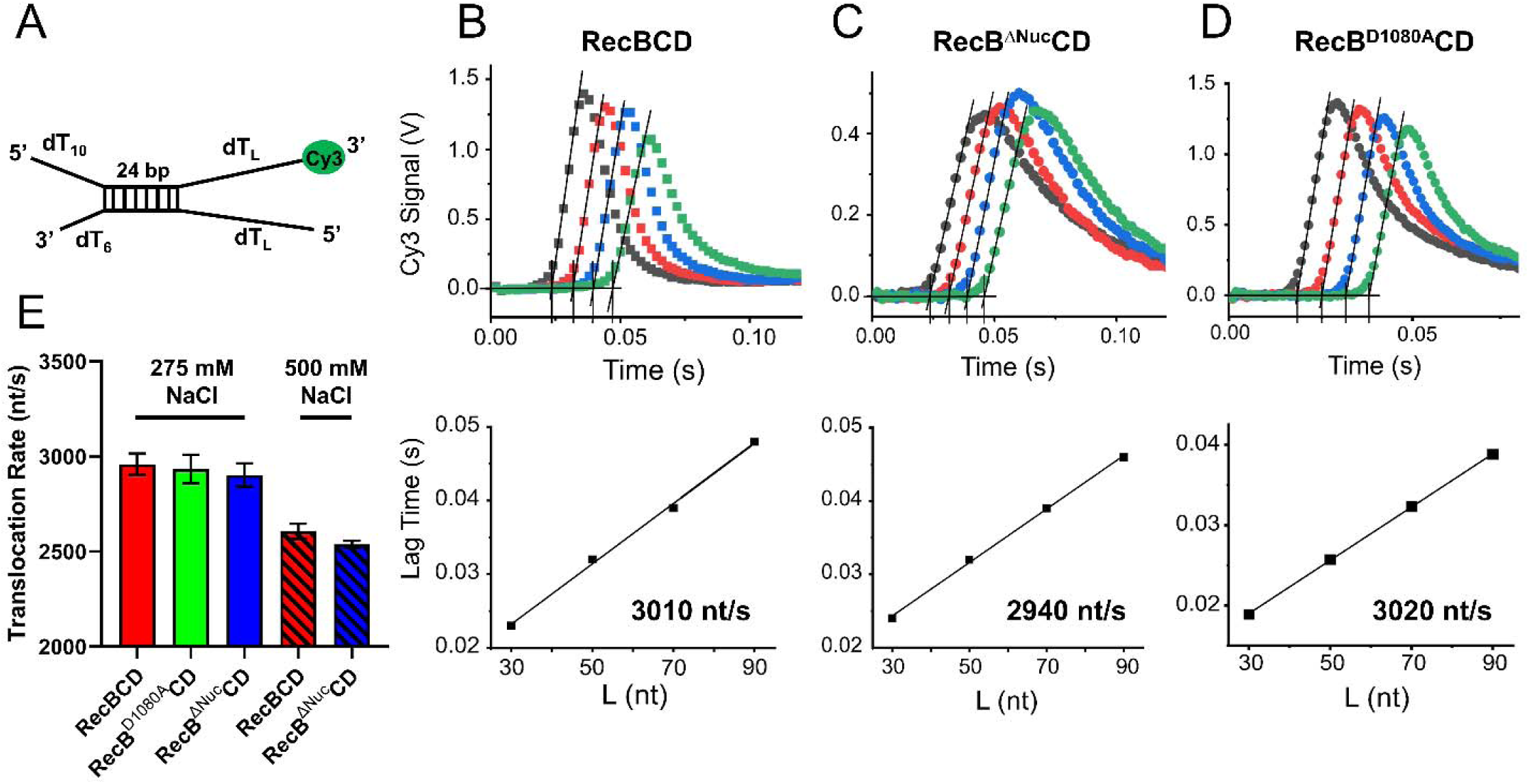
Stopped-flow time courses monitoring 5’ to 3’ ssDNA translocation of RecBCD, RecB^D1080A^CD, and RecB^ΔNuc^CD (M_275_ at 37°C, 5 mM ATP). **(A)** Pre-melted DNA substrate used to measure 5’ to 3’ ssDNA translocation. Representative DNA translocation time courses monitoring the enhancement of Cy3 fluorescence occurring when the helicase reaches the 5’ end of the ssDNA for **(B)** - RecBCD, **(C)** - RecB^D1080A^CD, and **(D)** - RecB^ΔNuc^CD. Time courses were obtained as a function of the 3’-ssDNA tail length (*L* = 30 nt (black); 50 nt (red); 70 nt (blue); 90 nt (green)). Lag times are plotted as a function of (L(nt)) below each data set and the macroscopic ssDNA translocation rate is estimated from the inverse of the slope of the best-fit line. **(E)** Average 5’ to 3’ ssDNA translocation rates for RecBCD, RecB^ΔNuc^CD, and RecB^D1080A^CD in buffer M_275_ (solid) and buffer M_500_ (striped) at 37°C and 5 mM ATP.

Translocation experiments were performed in buffer M_275_ and M_500_ at 37°C using a DNA hairpin trap (500 nM) to ensure single round conditions. These experiments were performed at [NaCl] ≥ 275 mM and 37°C and in excess DNA over enzyme to ensure that enzyme binding occurred exclusively at the 3’-dT_6_/5’-dT_10_ high affinity binding site. At lower [NaCl] and lower temperatures, the specificity of RecB^ΔNuc^CD for the high affinity 3’-dT_6_/5’-dT_10_ site is diminished such that RecB^ΔNuc^CD also binds to the labeled end of the single stranded tails. As described previously, this results in an initial dip in the Cy3 fluorescence signal, making quantitative analysis difficult [8].

The rates of ssDNA translocation were determined using a lag time analysis [7, 8, 51, 52] (see Materials and Methods). The lag time is determined as the time at the intersection of the initial linear increase in Cy3 fluorescence intersects the lag phase **(Fig. 6B-D, 7B, 7B-D)**. The rate of translocation is determined as the inverse slope of a plot of the lag time vs. ssDNA length **(Figures 6B-D, 7B-D)**. We attempted to fit the time courses to an n-step kinetic model; however, the 3’-5’ ssDNA translocation time courses for RecB^ΔNuc^CD could not be accurately fit by any models that we tried.

### Deletion of the nuclease domain does not affect the rate of ssDNA translocation

The results of ssDNA translocation experiments in buffer M_275_ at 37°C are shown for RecBCD **(Fig. 6B)**, RecB^ΔNuc^CD **(Fig. 6C)**, and RecB^D1080A^CD **(Fig.6 D)**. Four to five sets of experiments were performed for each enzyme (**Table S7)**. The average rate of 3’-5’ translocation for RecBCD is 2530 ± 40 nt/s, RecB^ΔNuc^CD is 2380 ± 110 nt/s, and for RecB^D1080A^CD is 2520 ± 50 nt/**s (**Fig. 6E, **Table 4)**. The average 5’-3’ translocation rate for RecBCD is 2960 ± 60 nt/s, RecB^ΔNuc^CD is 2900 ± 60 nt/s, and RecB^D1080A^CD is 2950 ± 85 nt/s **(Fig. 7E, Table 4)**. These results show that the rates of ssDNA translocation in both the 3’-5’ or 5’-3’ direction do not differ significantly (∼2%) for RecBCD vs. RecB^ΔNuc^CD. The rates of ssDNA translocation are significantly faster than the rates of DNA unwinding as noted previously [8]. We also note that the rate of ssDNA translocation by RecB^ΔNuc^CD in the 5’-3’ direction is slightly faster than in the 3’-5’ direction, which is also the case for RecBCD, consistent with previous reports [7, 8]. Translocation in the 5’ to 3’ direction is faster because both the RecD motor and the secondary translocase activity of the RecB motor contribute to the 5’-3’ translocation rate, whereas only the RecB motor contributes to translocation in the 3’-5’ direction [7, 8]. RecB^D1080A^CD, which retains the nuclease domain, but does not have nuclease activity, also shows the same ssDNA translocation rates as RecBCD and RecB^ΔNuc^CD (**Table 4**). We also examined the ssDNA translocation rates in buffer M_500_ at 37°C (500 mM NaCl). The ssDNA translocation rates in buffer M_500_ are the same for RecB^ΔNuc^CD and RecBCD, although slightly slower than the rates at 275 mM NaCl **(Table 4)**.

**Table 4.** Stopped-flow ssDNA translocation rates.

|  | Enzyme | 3' - 5' translocation rate (nt s<sup>-1</sup>) | 5' - 3' translocation rate (nt s<sup>-1</sup>) |
| --- | --- | --- | --- |
| M<sub>275</sub><br>37°C | BCD | 2530 ± 40 | 2960 ± 60 |
|  | B<sup>Δnuc</sup>CD | 2380 ± 110 | 2900 ± 60 |
|  | B<sup>D1080A</sup>CD | 2520 ± 50 | 2950 ± 85 |
| M<sub>500</sub><br>37°C | BCD | 2240 ± 25 | 2610 ± 40 |
|  | B<sup>Δnuc</sup>CD | 2190 ± 175 | 2540 ± 20 |

These results indicate that deletion of the nuclease domain slows the rate of DNA unwinding by 25%-45%, although the ssDNA translocation rate is unaffected. Therefore, the slower rate of DNA unwinding by RecB^ΔNuc^CD is not due to a slower rate of ssDNA translocation.

## Discussion

The changes in activity that occur upon recognition of a *chi*-site during DNA unwinding are some of the more interesting aspects of the RecBCD helicase. Upon *chi*-recognition, the enzyme pauses, but then the rate of DNA unwinding is reduced ∼two-fold and the nuclease activity changes from degrading both the 3’ and 5’-ended strands to degrading only the 5’ strand [25]. RecA is then loaded onto the 3’ strand possibly via an interaction with the nuclease domain [29]. These linkages between DNA unwinding and nuclease activity led us to examine whether the rate of DNA unwinding might be regulated by the state of the nuclease domain.

Using both ensemble stopped-flow and single DNA molecule optical tweezer approaches we show that the DNA unwinding (helicase) activity of RecBCD is regulated by its nuclease domain. Specifically, removal of the nuclease domain to form RecB^ΔNuc^CD slows the rate of DNA unwinding by as much as ∼50% under some conditions. However, the rates of ssDNA translocation are not affected by removal of the nuclease domain. This indicates that RecBCD helicase activity involves more than simple directional translocation along ssDNA. This conclusion is consistent with the observation that most monomeric SF1A ssDNA translocases show little to no helicase activity [53–58]. In addition, the binding affinity of RecB^ΔNuc^CD for a DNA end is higher than RecBCD [39]. This effect of removal of the nuclease domain is unexpected since in all RecBCD-DNA structures in which the nuclease domain is resolved [16–18], the nuclease domain is docked onto RecC at a site that is distant from the duplex DNA binding site, as shown in Figure 1. This suggests an allosteric effect of the nuclease domain on DNA binding and helicase activity.

The ensemble stopped-flow studies on short DNA substrates (**<=** 60 bp) showed a clear decrease in the rate of DNA unwinding (25-45%) by removal of the nuclease domain (RecB^ΔNuc^CD), whereas RecB^D1080A^CD, which contains the nuclease domain but has no nuclease activity, shows no change in the rate of DNA unwinding on short DNA substrates. This indicates that it is the nuclease domain, rather than nuclease activity that affects the DNA unwinding rate. The single molecule experiments, which use much longer DNA substrates (>19 kbp), also show a ∼two-fold decrease in the unwinding rates for RecB^ΔNuc^CD compared to RecBCD. However, on these longer DNA substrates, RecB^D1080A^CD shows an intermediate rate of DNA unwinding between RecBCD and RecB^ΔNuc^CD, suggesting that nuclease activity also can influence the rate of DNA unwinding.

The rate of initiation of DNA unwinding from a blunt ended DNA by RecB^ΔNuc^CD is also significantly slower than RecBCD and slower compared to its initiation from a 3’-dT_6_/5’-dT_10_ loading site. Efficient initiation of DNA unwinding from a DNA end has been proposed to require engaging both the RecB and RecD motors with each DNA strand, which requires DNA base pair (bp) melting and likely protein conformational changes[18, 59]. Single molecule studies demonstrated that the ATP-independent DNA melting by RecBCD from a DNA end is transient and dynamic, suggesting that even when RecBCD is pre-bound to DNA, both RecBCD motors must be engaged with ssDNA before DNA unwinding can be initiated [60, 61]. The slower initiation of DNA unwinding from a blunt DNA end by RecB^ΔNuc^CD suggests that the nuclease domain may influence DNA bp melting and motor engagement.

Recent cryo-EM structures of RecBCD bound to a blunt-ended DNA in the absence of ATP revealed two structural classes [18]. In one class, all domains of RecBCD are resolved, including the nuclease domain, which is observed in the same docked position as in previous crystal structures [16, 62]. In this structural class, at least 11 bp are melted from the DNA duplex end. In the second structural class, the nuclease domain is not visible, although it is present in the complex, and only four bp are melted from the duplex end. These results suggest that the nuclease domain can undock, but remain attached to the RecB motor domain via the 60 amino acid tether. It further suggests that the ability and extent of RecBCD to melt the DNA duplex may be regulated by the position of the nuclease domain. Cryo-EM structures of RecB^ΔNuc^CD bound to a blunt-duplex DNA end also show only four bp melted compared to up to 11 bp melted by RecBCD [18, 39].

Previous single DNA molecule optical tweezer studies of RecBCD helicase activity on DNA that does not contain *chi* sites have reported mostly continuous DNA unwinding, uninterrupted by significant pauses in the absence of a *chi* site [25, 28, 44, 47]. Significant pausing by RecBCD, by several seconds, was mainly observed upon *chi* site recognition by RecBCD [25, 28]. Furthermore, after *chi* recognition, the RecBCD unwinding rate was reduced by roughly two-fold [28]. This was attributed to a switching of the lead motors, wherein RecD is the faster motor prior to *chi* recognition, but RecB is the lead motor after *chi* recognition and the RecD motor might become disengaged [25]. However, our single DNA molecule optical tweezer studies show significant *chi-*independent pausing by RecBCD, as well as RecB^ΔNuc^CD and RecB^D1080A^CD. The duration of the pause times exhibited by RecBCD, RecB^ΔNuc^CD, or RecB^D1080A^CD are between ∼5.8 and 7.7 seconds. We note that the DNA unwinding duration times by RecBCD, RecB^ΔNuc^CD, or RecB^D1080A^CD are slightly shorter, 4.8-5.6 seconds, but are close to the pause duration times. However, in contrast to the *chi-*dependent pauses [25, 28], after which unwinding rates are reduced nearly two-fold, we do not observe a change in the distribution of DNA unwinding rates after these *chi-*independent pauses. It is important to note that the pauses and the unwinding events occurring after a pause are not from a new protein re-initiating unwinding, as both RecBC and RecBCD cannot initiate unwinding from long tailed substrates [7, 8]. In tethered particle single molecule experiments, Perkins et al. [63] also reported that RecBCD exhibited *chi*-independent pauses during DNA unwinding that were independent of both [ATP] and force.

In other single DNA molecule optical tweezer studies of RecBCD helicase activity, Liu et al. (2013) reported that even though the rates of unwinding by individual RecBCD enzymes varied substantially, an individual RecBCD did not pause while unwinding non *chi* site DNA [47]. Bimodal distributions of DNA unwinding rates were also observed by Liu et al. [47], which were attributed to a fast population in which both RecB and RecD motors were engaged with DNA and a slow population arising from either the RecB or RecD motors being disengaged. Liu et al. [47] further showed that transient pausing of RecBCD caused by depletion of Mg^2+^-ATP could change the DNA unwinding rates after re-introduction of the Mg^2+^-ATP. However, the DNA unwinding rate after the induced pause changed randomly from that found in the original population. We did not observe such bimodal activity in our single molecule studies for RecBCD or the two nuclease variants. Furthermore, the distribution of DNA unwinding rates after *chi-*independent pausing did not change significantly in our studies. Hence, the *chi-*dependent and *chi-*independent pausing show clear differences with respect to rate changes.

### Implications for the mechanism of RecBCD-catalyzed DNA unwinding

Despite much research, our understanding of the mechanism of DNA unwinding by RecBCD and most helicases remains incomplete [35]. A mechanical model based on structures of RecBCD bound to DNA suggests that DNA unwinding is a consequence of translocation by the canonical RecB and RecD motors along the respective single strands of DNA, which pulls the duplex DNA across a “wedge” or “pin” domain located within RecC, resulting in base pair separation [16]). This model requires that ssDNA translocation is tightly coupled to DNA melting/base pair separation, such that base pair separation only occurs when the motors translocate along their respective strands [16, 64]. This model predicts that the slower rate of dsDNA unwinding by RecB^ΔNuc^CD would also be accompanied by a slower rate of ssDNA translocation. However, we find that deletion of the nuclease domain does not affect the ssDNA translocation rates of either the RecB or RecD motors, although the DNA unwinding rate is reduced considerably. These observations are inconsistent with a simple “wedge” model of DNA unwinding.

An alternative model proposes that base pair melting occurs separately from ssDNA translocation [9, 11, 35]. In this model, the free energy of RecBCD binding to DNA is used to melt multiple DNA base pairs in an ATP-independent, Mg^2+^-dependent process [16, 18, 23, 24], forming ssDNA tracks along which the RecB and RecD motors translocate using ATP hydrolysis. Upon reaching the new ss/ds DNA junction, ATP hydrolysis resets the conformation of RecBCD, such that the binding free energy can again be used to melt the next set of multiple base pairs [35]. Previous biochemical studies have demonstrated that RecBCD, as well as RecBC, can melt multiple DNA base pairs upon binding to a DNA duplex end in an ATP-independent manner, i.e. using only its binding free energy [18, 23, 60, 61] [24, 40]. Structures of RecBCD bound to blunt-ended DNA in the absence of ATP show melting of 4-11 bp [16] [17, 18]. Further support for this model comes from Simon *et al*.[11], who demonstrated that duplex DNA unwinding can occur for at least 80 bp without ssDNA translocation by the canonical RecB and RecD motors. Interestingly, this activity also requires the nuclease domain but not nuclease activity [11]. As noted previously [8], and shown here as well, the rate of DNA unwinding is generally at least a factor of two slower than the rates of ssDNA translocation, also indicating that ssDNA translocation is not the rate limiting process for DNA unwinding, consistent with DNA unwinding being an active, rather than a passive process [35, 65–67]. We propose that the rate of DNA unwinding is limited by the rate of DNA base pair melting by RecBCD and that the slower rate of DNA unwinding by RecB^ΔNuc^CD, compared to RecBCD, may result from slower rates of the DNA melting step(s).

The observation that the DNA unwinding rate of RecB^ΔNuc^CD is 25-45% slower than RecBCD suggests that RecB^ΔNuc^CD may mimic the post chi state of RecBCD, in which the nuclease activity is altered and DNA unwinding is ∼two-fold slower [25]. This suggests the possibility that upon chi recognition, the nuclease domain is released from its docked position, although remaining covalently attached to the RecB motor domain through the flexible 60 amino acid linker. Undocking of the nuclease domain would make the previously occluded RecA binding site on the nuclease domain available to bind RecA and subsequently load RecA onto the resected 3’-ended ssDNA to form a RecA filament. However, recent studies [68] have concluded that the nuclease domain does not undock from the enzyme upon *chi-*recognition, but rather undergoes a *chi-*induced conformational change. It is possible that RecBCD, after such a conformational change of the nuclease domain, behaves similarly to RecB^ΔNuc^CD. Further studies will be needed to probe the dynamic nature of the RecB nuclease domain.

## Materials and Methods

### Buffers

Buffers were made with reagent grade chemicals and double-distilled water further deionized with a Milli-Q purification system (Millipore Corp., Bedford, MA) that were then filtered using 0.2 μm cellulose acetate filters (Corning, NY). Buffer C is 20 mM potassium phosphate, pH 6.8 at 4°C, 0.1 mM 2-mercaptoethanol, 0.1 mM EDTA, 10% (v/v) glycerol. Buffer M is 20 mM MOPS-KOH (pH 7.0), 5% (v/v) glycerol, 10 mM MgCl_2_, 1 mM 2-mercaptoethanol plus the indicated [NaCl]. A subscript denotes the [NaCl] in mM units (e.g., buffer M_275_ contains 275 mM NaCl). Stock MgCl_2_ concentrations were determined by refractive index using a Mark II refractometer (Leica Inc., Buffalo, NY).

Heparin (disodium salt (Sigma-Aldrich, St. Louis, MO)) stocks were prepared in Milli-Q water followed by extensive dialysis versus Milli-Q water (1 kDa cutoff (Spectrum Inc., Dominguez, CA)). Stock solutions were filtered and concentrations determined as described [69] and stored at 4°C.

ATP stocks were prepared by dissolving adenosine 5’-triphosphate sodium salt (Sigma, St. Louis, MO) in Milli-Q water and adjusting the pH to 7.0 using NaOH and concentrations were determined spectrophotometrically using ε_260_ = 1.54 × 10^4^ M^−1^cm^−1^ [70].

### Proteins

RecBCD was expressed from Plasmid pTRC99a in *E. coli* strain V2831 [71]. RecB^ΔNuc^CD, containing a deletion of amino acids 930-1180 of RecB, and RecB^D1080A^CD were expressed from plasmids pPB800 and pPB520, respectively, in *E. coli* strain V2601, a variant of V186 expressing the lacIq gene [72]. V2831 and V2601 contain deletions of the chromosomal *recB*, *recC*, and *recD* genes [39, 71]. Purification of RecBCD, RecB^D1080A^CD and RecB^ΔNuc^CD were as described [11, 33, 36, 39] with modifications [73]. We note that all RecBCD preps contained only hetero-trimeric complexes, with no significant hexameric complexes[74]. All proteins were ≥ 95% pure, assessed by sedimentation velocity analytical ultracentrifugation and SDS polyacrylamide gel electrophoresis and stored in buffer C at −80°C. Purified RecBCD, RecB^D1080A^CD, and RecB^ΔNuc^CD were dialyzed versus buffer M at 4°C before use and concentrations determined spectrophotometrically using ε_280_ = 4.5 × 10^5^ M^−1^ cm^−1^ (RecBCD and RecB^D1080A^CD) and ε_280_ = 4.11 × 10^5^ M^−1^ cm^−1^ (RecB^ΔNuc^CD) [11, 36]. Bovine serum albumin (BSA) (Sigma St. Louis, MO) was dialyzed vs. buffer M and concentrations determined spectrophotometrically using the extinction coefficient, ε_280_ = 4.38 × 10^4^ M^−1^cm^−1^ [40, 75]. SSB protein was a gift from Alex Kozlov and was purified as described [76].

### DNA

Oligodeoxynucleotides were synthesized using a MerMade 4 synthesizer (Bioautomation, Plano, TX) with phosphoramidites (Glen Research, Sterling, VA) and purified as described [77]. DNA concentrations were determined as described [11, 78] by digesting with phosphodiesterase I (Worthington, Lakewood, NJ) in PBS buffer at 25°C to form a mixture of mononucleotides and determining the absorbance of the mixture of mononucleotides at 260 nm. The extinction coefficient at 260 nm was calculated as the sum of individual mononucleotides (ε_260_ = 15,340 M^−1^ cm^−1^ for AMP, ε_260_ = 7600 M^−1^ cm^−1^ for CMP, ε_260_ = 12,160 M^−1^ cm^−1^ for GMP, ε_260_ = 8700 M^−1^cm^−1^ for TMP) and ε_260_ = 4930 M^−1^ cm^−1^ for Cy3 and ε_260_ = 10000 M^−1^ cm^−1^ for Cy5 (Glen Research) based on the sequence of each DNA strand **(Supplementary Table S1)**. Equal concentrations of Cy3-labeled ssDNA, Cy5-labeled ssDNA and unlabeled ssDNA were mixed to form the DNA unwinding substrates. For the ssDNA translocation substrates, the Cy3 labeled and unlabeled strands were mixed in equal concentrations, heated to 95°C for 5 minutes and allowed to cool slowly to 25°C.

### Fluorescence stopped-flow experiments

Fluorescence stopped-flow experiments were performed at 37°C in buffer M at the indicated [NaCl] using an Applied Photophysics SX.18MV instrument (Applied Photophysics Ltd., Leatherhead, UK). DNA was preincubated with enzyme in buffer M containing 6 μM BSA for 5 min on ice and then loaded into one syringe of the stopped-flow instrument. The other syringe contained a solution of ATP, 6 μM BSA, and protein trap to ensure single round conditions. Solutions were incubated for 3 min at 37°C in the instrument before 1:1 mixing.

For DNA unwinding, the final conditions after mixing were 25 nM RecBCD (or variant), 20 nM DNA, 6 μM BSA, 5 mM ATP, protein trap as noted. Cy3 fluorescence was excited using a 505 nm LED (Applied Photophysics Ltd.). Cy3 and Cy5 fluorescence were monitored simultaneously using separate photomultipliers containing a 570 nm interference filter and a >665 nm-cutoff filter (Oriel Corp., Stradford, CT), respectively.

For ssDNA translocation, the final conditions after mixing were 37.5 nM RecBCD (or variant), 50 nM DNA, 6 μM BSA, 5 mM ATP, 500 nM hairpin trap. Cy3 fluorophore was excited using a 505 nm LED (Applied Photophysics Ltd.) and its emission was monitored using a >570 nm-cutoff filter (Oriel Corp., Stradford, CT). For dsDNA unwinding and ssDNA translocation, the kinetic traces shown represent the average of at least 8 individual measurements. Experiments were done in triplicate, except for the RecB^D1080A^CD unwinding experiments, which were performed once. No differences in results were noted for experiments performed using protein from multiple preparations.

Heparin was used as the protein trap for experiments done at 30 mM NaCl, but at higher [NaCl] (275 mM and 500 mM), heparin is not an effective trap and a 16 bp DNA hairpin with a 3’-dT_6_/5’-dT_10_ end was used as the trap. ssDNA translocation experiments were performed at higher [NaCl] to ensure initiation of DNA unwinding only from the 3’-dT_6_/5’-dT site. The final trap concentrations used were: RecBCD and RecB^ΔNuc^CD dsDNA unwinding experiments in buffer M_30_ with 4 and 2 mg/ml heparin, respectively; RecBCD and RecB^ΔNuc^CD dsDNA unwinding and ssDNA translocation experiments in buffer M_275_ and buffer M_500_ – 500 nM DNA hairpin trap.

### Analysis of stopped-flow DNA unwinding time courses

DNA unwinding time courses were fit globally to the n-step kinetic model in Scheme 1 [36, 37] using MENOTR, a hybrid multi-start genetic and non-linear least squares (NLLS) algorithm [38]. Time courses monitoring the Cy5 signal, were analyzed out to 0.5 s for RecBCD and RecB^D1080A^CD and 2 s for RecB^ΔNuc^CD to capture the signal plateau.

In Scheme 1, the DNA is unwound via *n* repeated steps with step rate constant, *k_U_*, and a step size of *m* bp. The pre-bound helicase-DNA complexes start as a mixture of two states: a non-productive state, (*RD*)*_NP_*, and a productive state, (*RD*)*_P_*. The fraction of complexes in the productive state, (*RD*)*_P_*, is given by *x* (Eq. 1), and the fraction in the non-productive state, (*RD*)*_NP_*, is given by (1-*x*). Unwinding initiates from the (*RD*)*_P_* state, hence the non-productive complexes (*RD*)*_NP_* must first isomerize with rate constant *k_NP_* to form (*RD*)*_P_*before unwinding can initiate. Productive (*RD*)*_P_* complexes must also proceed through a number (*h*) of kinetic steps with rate constant, *k_C_*, before unwinding can occur via the repeated unwinding steps with rate constant, *k_U_*. In the global NLLS analysis, *k_U_*, *k_C_*, *h*, *k_NP_*, and *x* were constrained as global parameters (i.e., the same for each time course). The number of *k_C_* steps, *h*, and the fraction of initially productive complexes, *x*, have been shown to be independent of the DNA length, *L*, that is unwound [36, 40, 79]. The number of *k_U_* steps, *n*, and the amplitude terms, *A* and *B*, were determined as fitted parameters for each DNA length, *L*. The macroscopic unwinding rate is the product, *mk_U_*, and is better constrained than the individual values of *k_U_* and *m*, as these parameters are highly correlated.

Fitting of the time courses to Scheme 1 (Eq. (2) or (3)) was performed to obtain the time-dependent formation of ssDNA*, f_ss_(t),* plus the formation of the last intermediate, *I_n-1_*, as the inverse Laplace transform of *F_ss_(s)* using numerical methods as described [33, 36, 37]. *F_ss_(s)* is the Laplace transform of *f_ss_(t)*, where L ^−1^ is the inverse Laplace transform operator with *s* as the Laplace variable, *A* is the amplitude of the fluorescence change upon ssDNA formation, *B* is an amplitude term that accounts for the fluorescence enhancement due to formation of the last intermediate before complete DNA unwinding *I_n-1_*. The number of unwinding steps, *n*, is related to *L* and *m* by Eq. (4).

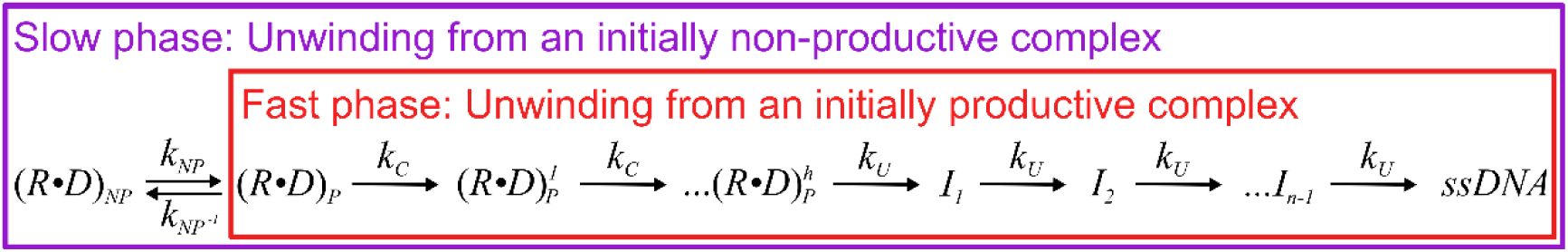
Scheme 1.

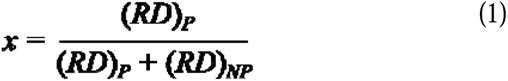

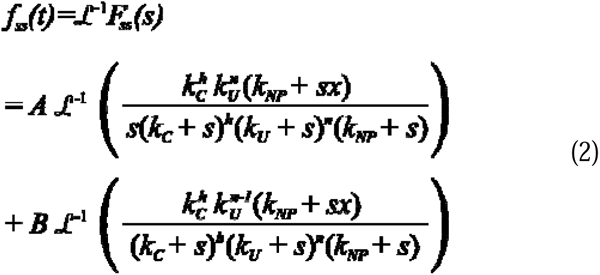

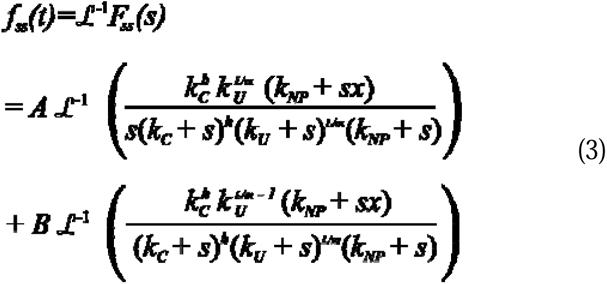

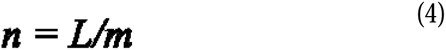

### Analysis of ssDNA translocation time courses

The rate of ssDNA translocation was calculated from the dependence of the “lag-time”, on ssDNA length as described [7, 8, 51]. The “lag time” was determined as the time at the intersection of the linear increase of the Cy3 fluorescence with the initial Cy3 fluorescence and reflects the time for RecBCD (or variant) to unwind the 24 bp duplex DNA and translocate to the Cy3 fluorophore at the end of the ssDNA, whereupon an increase in Cy3 fluorescence occurs[50, 80]. The lag time is determined for a series of DNA substrates differing in ssDNA length, and the inverse slope of a plot of ssDNA length versus lag time yields the ssDNA translocation rate [7, 8, 51]. Because the 3′-dT_6_/5’-dT_10_ tail and the duplex length (24 bp) is constant for all DNA substrates, the slope of the plot is dependent only on the length of the ssDNA. Reported translocation rates are the average from all experiments and the uncertainties are given as the standard deviation.

### Single DNA molecule LUMICKS C-Trap experiments

Single molecule DNA unwinding experiments were performed with a LUMICKS C-Trap controlled with Bluelake^TM^ (v2.0) software. The combined optical tweezer and confocal scanning microscope is furnished with a μ-Flux™ Microfluidics System (LUMICKS) attached to a flow cell containing 5 channels (C1, LUMICKS). The LUMICKS C-Trap was passivated prior to performing experiments by flowing PBS Buffer (0.5 mL at 1.6 bar) through the syringes, lines, and flow cell, followed by flowing 0.5% (w/v) pluronic acid (0.5 mL at 1.6 bar, and Pluronic® F-127 P2443, Sigma) and then 0.1% (w/v) BSA (0.5 mL at 1.6 bar, A-6793, Sigma). The flow cell, syringes, and tubing were passivated overnight in the PBS Buffer containing 0.1% (w/v) BSA. The passivation procedure ended by flowing PBS Buffer (0.5 mL at 1.6 bar) through the system before beginning experiments the next day.

The DNA unwinding experiments utilized a 19,435 base pair duplex DNA with a blunt-end and a 3’ biotinylated end on the opposite end which can be attached to a streptavidin-coated polystyrene bead. The DNA was generated by treating 40ng of 20,452-bp biotinylated bacteriophage λ-DNA (SKU00014, LUMICKS) with 20U of SmaI (R0141S, New England BioLabs) for 25 minutes at 25°C, followed by an inactivation step (65°C for 30 min.). Aliquots of the DNA were placed at −20°C for long-term storage.

The experimental procedure is summarized in **Supplementary Figure S3**. A streptavidin-coated polystyrene bead (4.34 μm, 0.5% (w/v), Cat No.: SVP-40-5, Spherotech Inc.) with a single 19,435 bp DNA tether bound to its surface was captured with Trap 2 in Channel 1. Roughly one DNA tether was bound per bead mixing a 1:450 dilution of streptavidin-coated polystyrene beads and 1 ng of premade DNA tether in PBS Buffer and 250 nM Sytox Orange (Invitrogen, S11368). A bead and single DNA tether held within Trap 2 were moved to Channel 2 containing Buffer M_30_ supplemented with Sytox Orange (250 nM). After initiating imaging, the bead and tether were moved into Channel 3, containing Buffer M_30_, 250 nM Sytox Orange, 2 mM ATP, and 100 nM of RecBCD, RecB^ΔNuc^CD or RecB^D1080A^CD. Experiments in the presence of *E. coli* SSB were performed by supplementing the Channel 3 solution with 124 nM of SSB. All single molecule DNA unwinding experiments were performed at 28°C.

Confocal scans of the DNA tether during unwinding were acquired using a 532 nm laser at 5% power. The full 7.53 x 1.82 µm imaging area was scanned in 489.6 ms increments, with a line time of 27.2 ms, and a pixel size of 100 nm. Movies were converted to TIFF stacks which were subsequently analyzed by hand using the measurement tool in ImageJ. The measurements from the trajectories were normalized to 19,435 bp and plotted in GraphPad Prism from which the data were analyzed to obtain DNA unwinding rates. Each of the single molecule DNA experiments was performed under flow by placing the μ-Flux™ Microfluidics System pressure at 0.18 bar. The flow rate was determined in separate experiments to be 1402 ± 46 μm/s (mean ± SD (n=10)) by imaging the movement of a 4.34 μm polystyrene bead released from a trap when the microfluidics system pressure was held at 0.18 bar in PBS Buffer at 28°C.

## Acknowledgements

We thank members of the Lohman lab and Roberto Galletto for helpful discussions, Thang Ho for synthesis and purification of the oligodeoxynucleotides, Alex Kozlov for providing purified SSB protein and Zach Ingram and Aaron Lucius for help with implementing MENOTR. NF thanks Drake Jensen for helpful discussions. This research was supported in part by the NIH (R35 GM136632 to TML). The LUMICKS C-Trap G2 was purchased with support from NIH Instrumentation grant S10OD030315.

