## Supplemental Material for "*E. coli* RecBCD Nuclease Domain Regulates Helicase Activity but not Single Stranded DNA Translocase Activity"

^2^Current address: Department of Chemistry, Washington University in St. Louis, St. Louis, MO 63130

^3^Current address: KBI Biopharma Inc., Boulder, CO

**Supplemental Tables**

**Table S1. Sequences for dsDNA unwinding and ssDNA translocation substrates.**

T_6_T_10_ Hairpin trap:

5’ (dT)_10_TCG AAG TAG ACA GAT CCT AGT GCA GG TTT TCC TGC ACT AGG ATC TGT CTA CTT CG (dT)_6_ 3’

Unwinding Substrates:

Cy3 labeled unwinding substrates:

*L* (nt)	DNA sequence

25 (25) 5’ ((dT)_10_) TGT GTC ACA GTT ACT CAG ACT TGA A Cy3-3’

30 (30) 5’ ((dT)_10_) TGT GTC ACA GTT ACT CAG ACT TGA ATG TCA Cy3-3’

37 (37) 5’ ((dT)_10_) TGT GTC ACA GTT ACT CAG ACT TGA ATG TCA TGT CTC G Cy3 -3’

40 (40) 5’ ((dT)_10_) TGT GTC ACA GTT ACT CAG ACT TGA ATG TCA TGT CTC GAA A Cy3 -3’

43 (43) 5’ ((dT)_10_) TGT GTC ACA GTT ACT CAG ACT TGA ATG TCA TGT CTC GAA ATC A Cy3 -3’

48 (48) 5’ ((dT)_10_) TGT GTC ACA GTT ACT CAG ACT TGA ATG TCA TGT CTC GAA ATC ATC CAT Cy3 -3’

50 (50) 5’ ((dT)_10_) TGT GTC ACA GTT ACT CAG ACT TGA ATG TCA TGT CTC GAA ATC ATC CAT GT Cy3 -3’

53 (53) 5’ ((dT)_10_) TGT GTC ACA GTT ACT CAG ACT TGA ATG TCA TGT CTC GAA ATC ATC CAT GTG CT Cy3 -3’

60 (60) 5’ ((dT)_10_) TGT GTC ACA GTT ACT CAG ACT TGA ATG TCA TGT CTC GAA ATC ATC CAT GTG CTA GAG ACA Cy3 -3’

Cy5 labeled unwinding substrates:

*L* (nt)	DNA sequence

25 (55) 5’ Cy5 TG TCA TGT CTC GAA ATC ATC CAT GTG CTA GAG ACA TCA TCT GTA ATG AAG TAG AC 3’

30 (50) 5’ Cy5 TGT CTC GAA ATC ATC CAT GTG CTA GAG ACA TCA TCT GTA ATG AAG TAG AC 3’

37 (43) 5’ Cy5 AA ATC ATC CAT GTG CTA GAG ACA TCA TCT GTA ATG AAG TAG AC 3’

40 (40) 5’ Cy5 TC ATC CAT GTG CTA GAG ACA TCA TCT GTA ATG AAG TAG AC 3’

43 (37) 5’ Cy5 TC CAT GTG CTA GAG ACA TCA TCT GTA ATG AAG TAG AC 3’

48 (32) 5’ Cy5 GTG CTA GAG ACA TCA TCT GTA ATG AAG TAG AC 3’

50 (30) 5’ Cy5 G CTA GAG ACA TCA TCT GTA ATG AAG TAG AC 3’

53 (27) 5’ Cy5 TA GAG ACA TCA TCT GTA ATG AAG TAG AC 3’

60 (20) 5’ Cy5 TCA TCT GTA ATG AAG TAG AC 3’

Unwinding Template strand (underlined sequence forms 15 bp hairpin with 4 nt loop)

5’ AGA TCC TAG TGC AGG TTT TCC TGC ACT AGG ATC TGT CTA CTT CAT TAC AGA TGA TGT CTC TAG CAC ATG GAT GAT TTC GAG ACA TGA CAT TCA AGT CTG AGT AAC TGT GAC ACA ((dT)_6_) 3’

Translocation 3’ – 5’ substrates:

Top strands:

5’-Cy3 (dT)_30_ CTG CAG CTA GCT CAG GAG CCA TGG (dT)_6_ – 3’

5’-Cy3 (dT)_50_ CTG CAG CTA GCT CAG GAG CCA TGG (dT)_6_ – 3’

5’-Cy3 (dT)_70_ CTG CAG CTA GCT CAG GAG CCA TGG (dT)_6_ – 3’

5’-Cy3 (dT)_90_ CTG CAG CTA GCT CAG GAG CCA TGG (dT)_6_ – 3’

Bottom Strands:

5´- (dT)_10_ CCA TGG CTC CTG AGC TAG CTG CAG (dT)_30_ -3´

5´- (dT)_10_ CCA TGG CTC CTG AGC TAG CTG CAG (dT)_50_ -3’

5´- (dT)_10_ CCA TGG CTC CTG AGC TAG CTG CAG (dT)_70_ -3’

5´- (dT)_10_ CCA TGG CTC CTG AGC TAG CTG CAG (dT)_90_ -3’

Translocation 5’-3’ substrates:

Top strands:

5´- (dT)_10_ CCA TGG CTC CTG AGC TAG CTG CAG (dT)_30_ Cy3-T-3´

5´- (dT)_10_ CCA TGG CTC CTG AGC TAG CTG CAG (dT)_50_ Cy3-T-3´

5´- (dT)_10_ CCA TGG CTC CTG AGC TAG CTG CAG (dT)_70_ Cy3-T-3´

5´- (dT)_10_ CCA TGG CTC CTG AGC TAG CTG CAG (dT)_90_ Cy3-T-3´

Bottom strands:

5’- (dT)_30_ CTG CAG CTA GCT CAG GAG CCA TGG (dT)_6_ – 3’

5’- (dT)_50_ CTG CAG CTA GCT CAG GAG CCA TGG (dT)_6_ – 3’

5’- (dT)_70_ CTG CAG CTA GCT CAG GAG CCA TGG (dT)_6_ – 3’

5’- (dT)_90_ CTG CAG CTA GCT CAG GAG CCA TGG (dT)_6_ – 3’

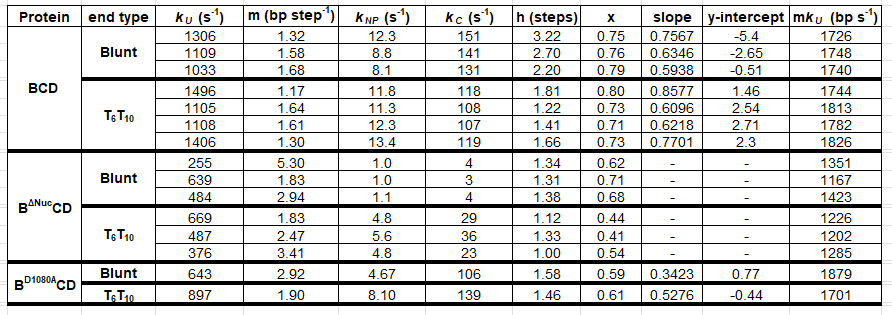
**Table S2. DNA Unwinding rate constants for all data sets collected in Buffer M_30_.**

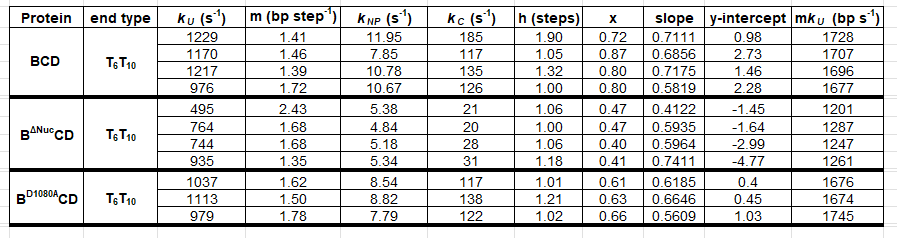
**Table S3. DNA unwinding rate constants for all data sets collected in Buffer M_275_.**

**Table S4. DNA unwinding rate constants for all data sets collected in Buffer M_500_.**

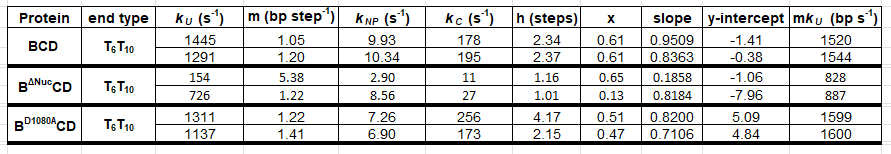

**Table S5: DNA unwinding rates for RecBCD, RecB^D1080A^CD, and RecB^∆Nuc^CD determined by single-molecule (LUMICKS C-TRAP) experiments.**

| **RecBCD or Variant** | **rates of unwinding** | **std. dev.** | **# of unwinding events** | **# of trajectories** |
| --- | --- | --- | --- | --- |
| **RecBCD*** | 1612 | 572 | 84 | 84 |
| **RecB^D1080A^CD*** | 1084 | 503 | 26 | 26 |
| **RecB^∆Nuc^CD*** | 723 | 420 | 42 | 42 |
| **RecBCD** | 1615 | 721 | 218 | 144 |
| **RecB^D1080A^CD** | 1117 | 596 | 95 | 55 |
| **RecB^∆Nuc^CD** | 883 | 658 | 109 | 72 |
| **RecBCD + SSB** | 2035 | 788 | 106 | 82 |
| **RecB^D1080A^CD + SSB** | 1270 | 707 | 101 | 57 |
| **RecB^∆Nuc^CD + SSB** | 1033 | 711 | 136 | 78 |

*** Trajectories containing pauses were omitted**

**Table S6: DNA unwinding processivities for RecBCD, RecB^D1080A^CD, and RecB^∆Nuc^CD with and without *E. coli* SSB protein.**

| **RecBCD or Variant** | **Processivity** | **Basepairs Unwound** |
| --- | --- | --- |
| **RecBCD** | 0.99995 | 21,165 |
| **RecB^D1080A^CD** | 0.99993 | 14,192 |
| **RecB^∆Nuc^CD** | 0.99992 | 12,823 |
| **RecBCD + SSB** | 0.99996 | 23,386 |
| **RecB^D1080A^CD + SSB** | 0.99994 | 17,126 |
| **RecB^∆Nuc^CD + SSB** | 0.99993 | 15,366 |

**Processivity is P = e^-0.693/x^, x is when ½ of the motors have ceased unwinding**

**Table S7. ssDNA translocation rates for all data sets.**

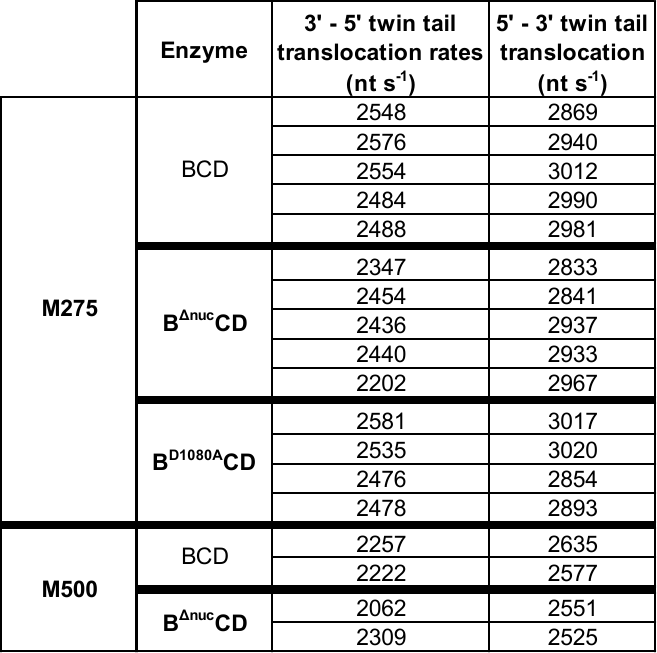

**Supplemental Data**

A

B

C

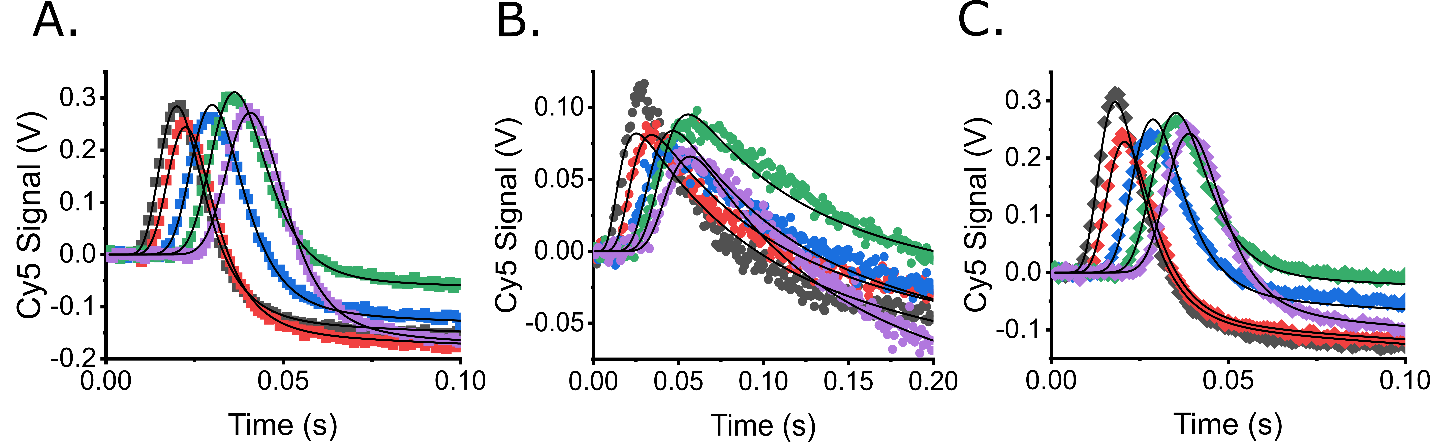

**Figure S1.** Representative unwinding time courses and simulated fits to Scheme 1 for DNA unwinding from a 3’-dT_6_/5’-dT_10_ DNA end in buffer M_275_ at 37ºC for **A)** RecBCD, **B)** RecB^ΔNuc^CD, and **C)** RecB^D1080A^CD. Colored symbols are data and smooth black lines are global fits to Scheme 1. Lengths shown are as follows: *L* = 25 bp (black), 30 bp (red), 40 bp (blue), 50 bp (green) and 60 bp (purple). Average best fit parameters are indicated in the text and Table 2 and best fit parameters for each data set are in Supplementary Table S3.

A

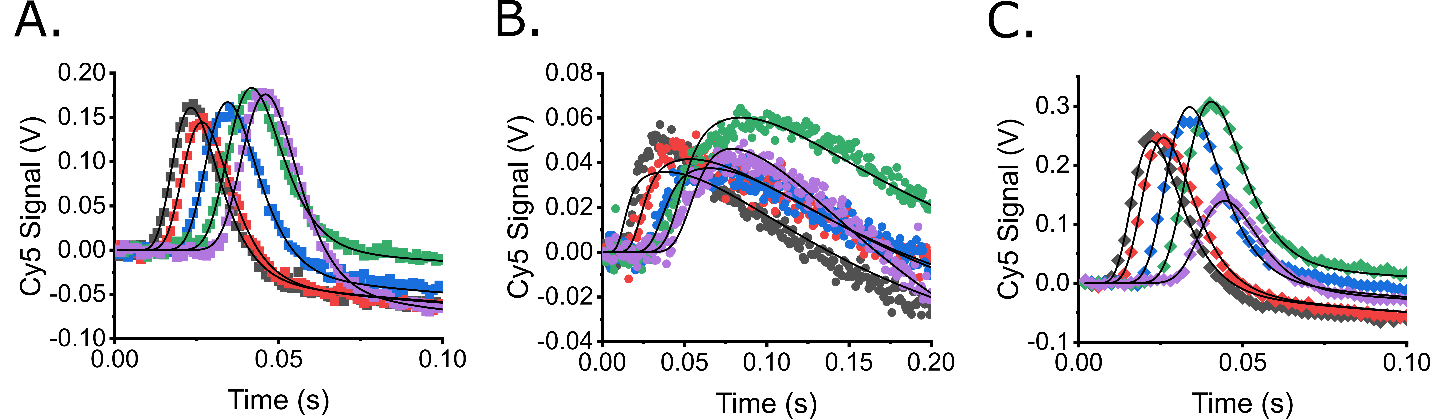

B

C

**Figure S2.** Representative unwinding time courses and simulated fits to Scheme 1 from a 3’-dT_6_/5’-dT_10_ DNA end in buffer M_500_ at 37ºC for **A)** RecBCD, **B)** RecB^ΔNuc^CD, and **C)** RecB^D1080A^CD. Colored symbols are data and smooth black lines are global fits to Scheme 1. Lengths shown are as follows: *L* = 25 bp (black), 30 bp (red), 40 bp (blue), 50 bp (green) and 60 bp (purple). Average best fit parameters are indicated in the text and Table 3 and best fit parameters for each data set are in Supplementary Table S4.

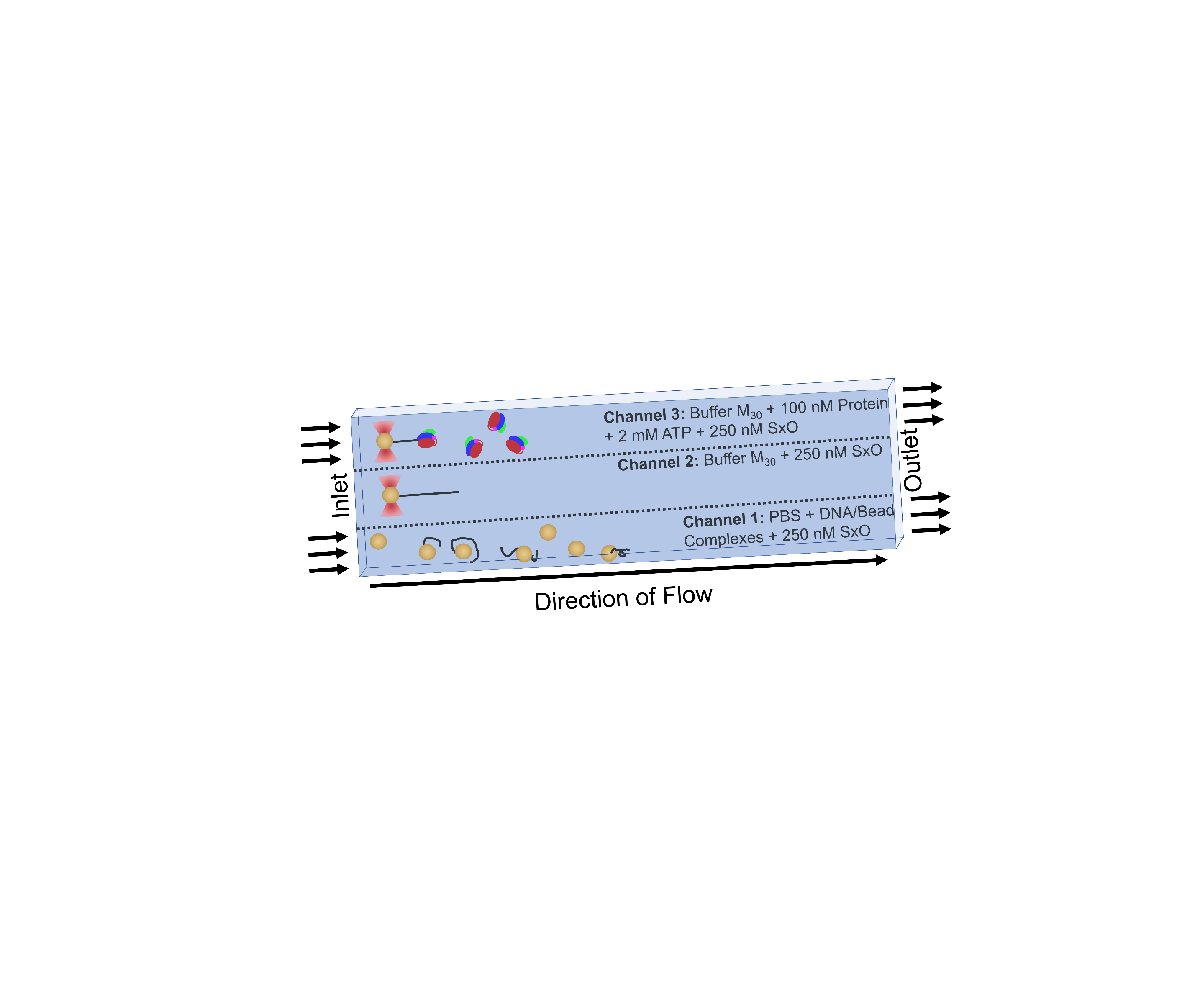

**Figure S3. Flow cell layout for LUMICKS C-trap single molecule unwinding experiments.** The three channels are separated into three lanes (dotted lines) by laminar flow. Polystyrene beads complexed with DNA were selected for single tethers in Channel 1. The timelapse recordings were started in Channel 2 before moving the bead and DNA assembly into Channel 3 that also contains ATP. Once the bead and DNA tether assembly were moved into channel 3, a single helicase initiates DNA unwinding from the free blunt-end of the DNA. Sytox Orange (250 nM) was present in each channel to confirm a single DNA tether is bound during the set-up and imaging.

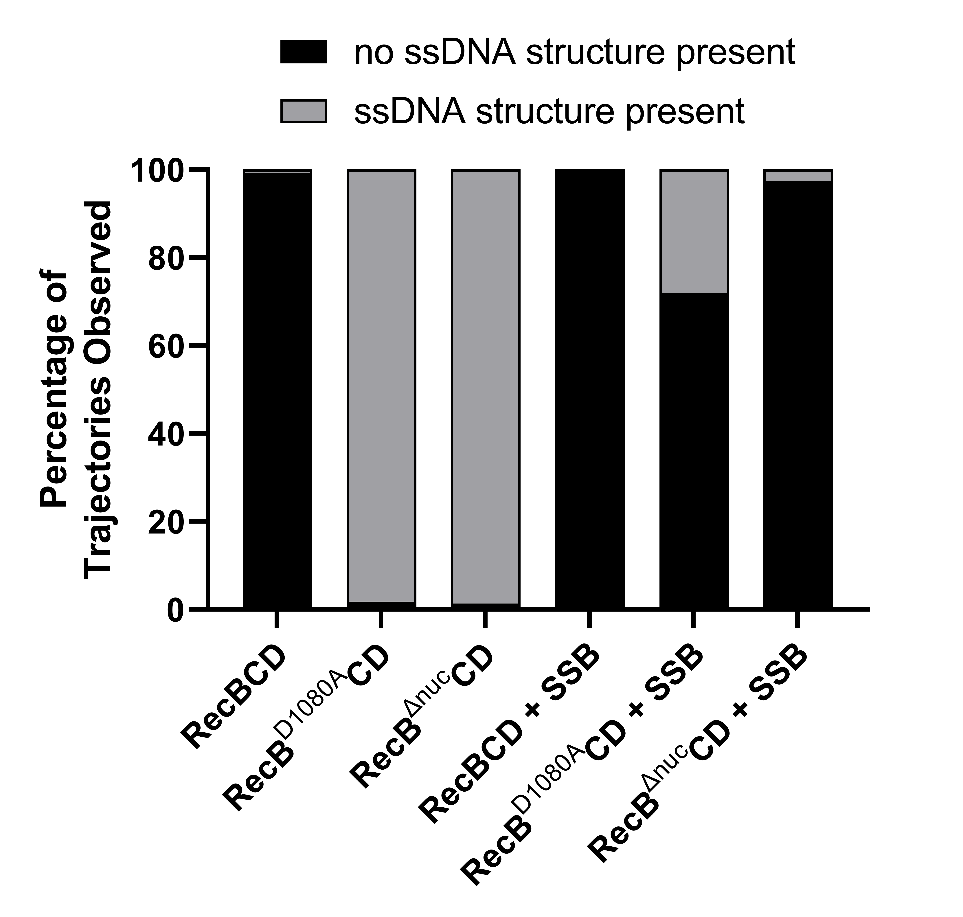

**Figure S4.** Quantification of DNA trajectories showing fluorescent spots. The percentage of trajectories exhibiting the formation of ssDNA structures during unwinding across all groups measured (gray = percentage of trajectories that form DNA structures, black = percentage of trajectories that do not form secondary DNA structures. % of trajectories that do not form secondary DNA structures, (n = number of trajectories) RecBCD: 99.3 %, n = 144; RecB^ΔNuc^CD: 0 %, n = 72; RecB^D1080A^CD: 0 %, n = 55; RecBCD + SSB: 100 %, n = 82; RecB^ΔNuc^CD + SSB: 97.4%, n = 78; RecB^D1080A^CD + SSB: 71.9 %, n = 57.

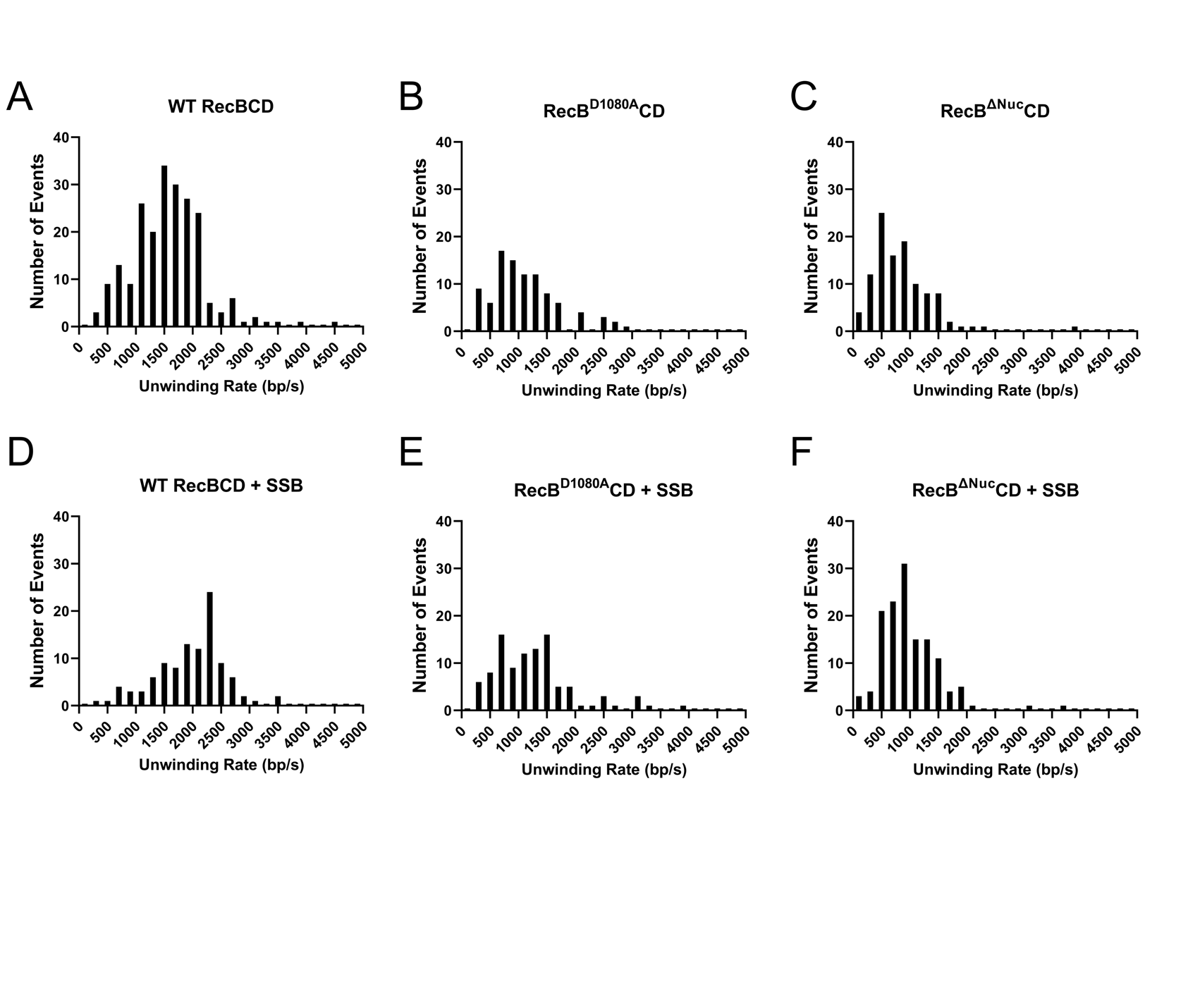

**Figure S5. DNA unwinding rate distributions of RecBCD, RecB^D1080A^CD, and RecB^∆Nuc^CD in the absence and presence of 124 nM *E. coli* SSB. A-F)** Distributions presented here include data from fitting every unwinding event within a trajectory. **A)** WT RecBCD (n = 218), **B)** RecB^D1080A^CD (n = 95), **C)** and RecB^∆Nuc^CD (n = 109), **D)** WT RecBCD (n = 106), **E)** RecB^D1080A^CD (n = 101), **F)** RecB^∆Nuc^CD (n = 136).

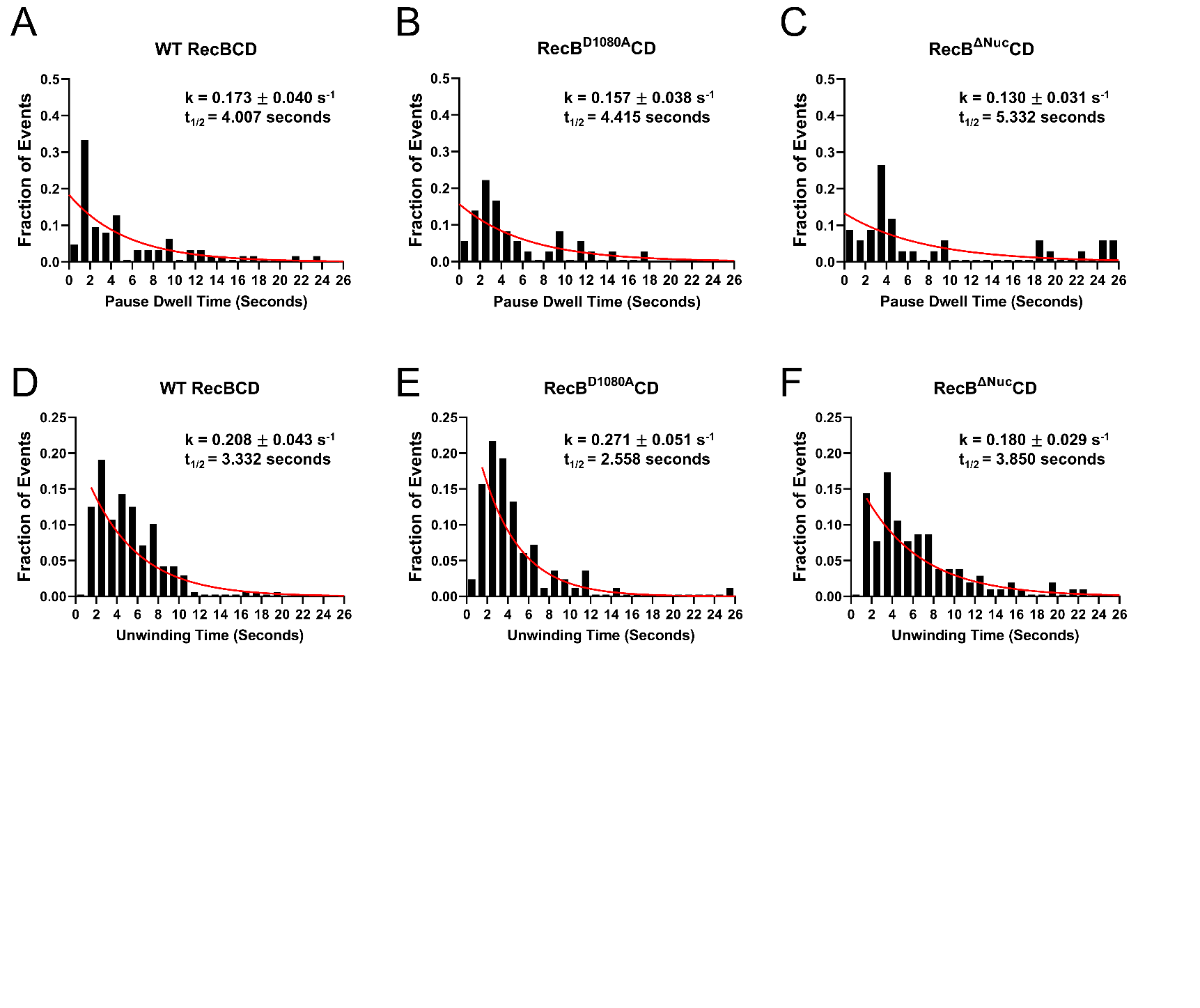

**Figure S6. Pause dwell times and DNA unwinding times from LUMICKS C-TRAP experiments. A-F)** Pause dwell times and DNA unwinding times for RecBCD, RecB^D1080A^CD, and RecB^∆Nuc^CD were quantified fit to an exponential probability density function in red (y=ke^-kx^). Rates (k) for each distribution are presented on the plot with the standard deviation of the fit. **A)** WT RecBCD (n=65), **B)** RecB^D1080A^CD (n=36), **C)** and RecB^∆Nuc^CD (n=32), **D)** WT RecBCD (n=168) **E)** RecB^D1080A^CD (n=83) **F)** RecB^∆Nuc^CD (n=104).

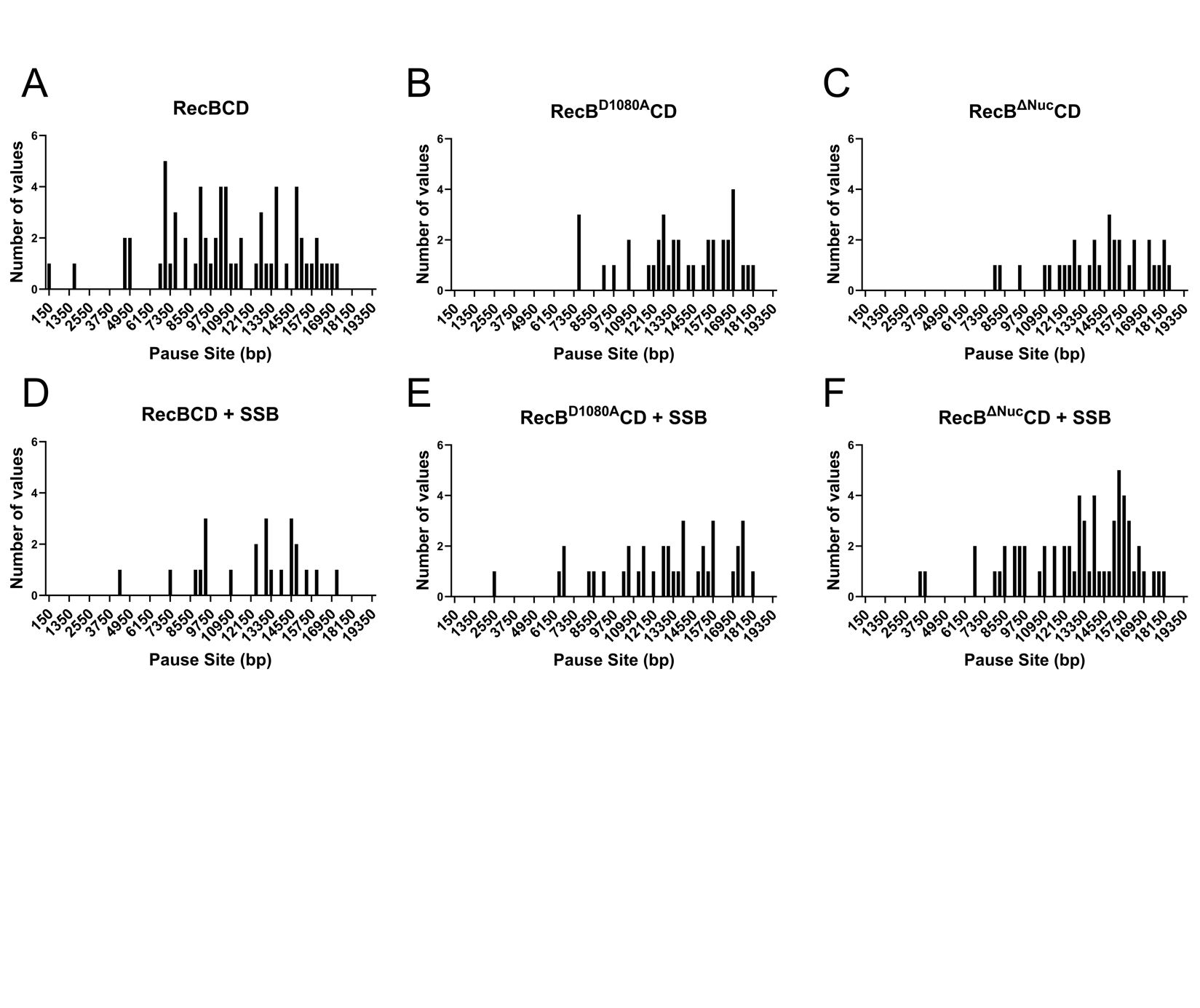

**Figure S7. DNA unwinding pauses occur randomly. A-F)** The locations of RecBCD, RecB^D1080A^CD, and RecB^∆Nuc^CD in both absence and presence of 124 nM *E. coli* SSB are plotted here and are shown to occur at random locations along the DNA. **A)** WT RecBCD (n = 65), **B)** RecB^D1080A^CD (n = 37), **C)** and RecB^∆Nuc^CD (n = 32), **D)** WT RecBCD + SSB (n = 23) **E)** RecB^D1080A^CD + SSB (n = 38) **F)** RecB^∆Nuc^CD + SSB (n = 61).

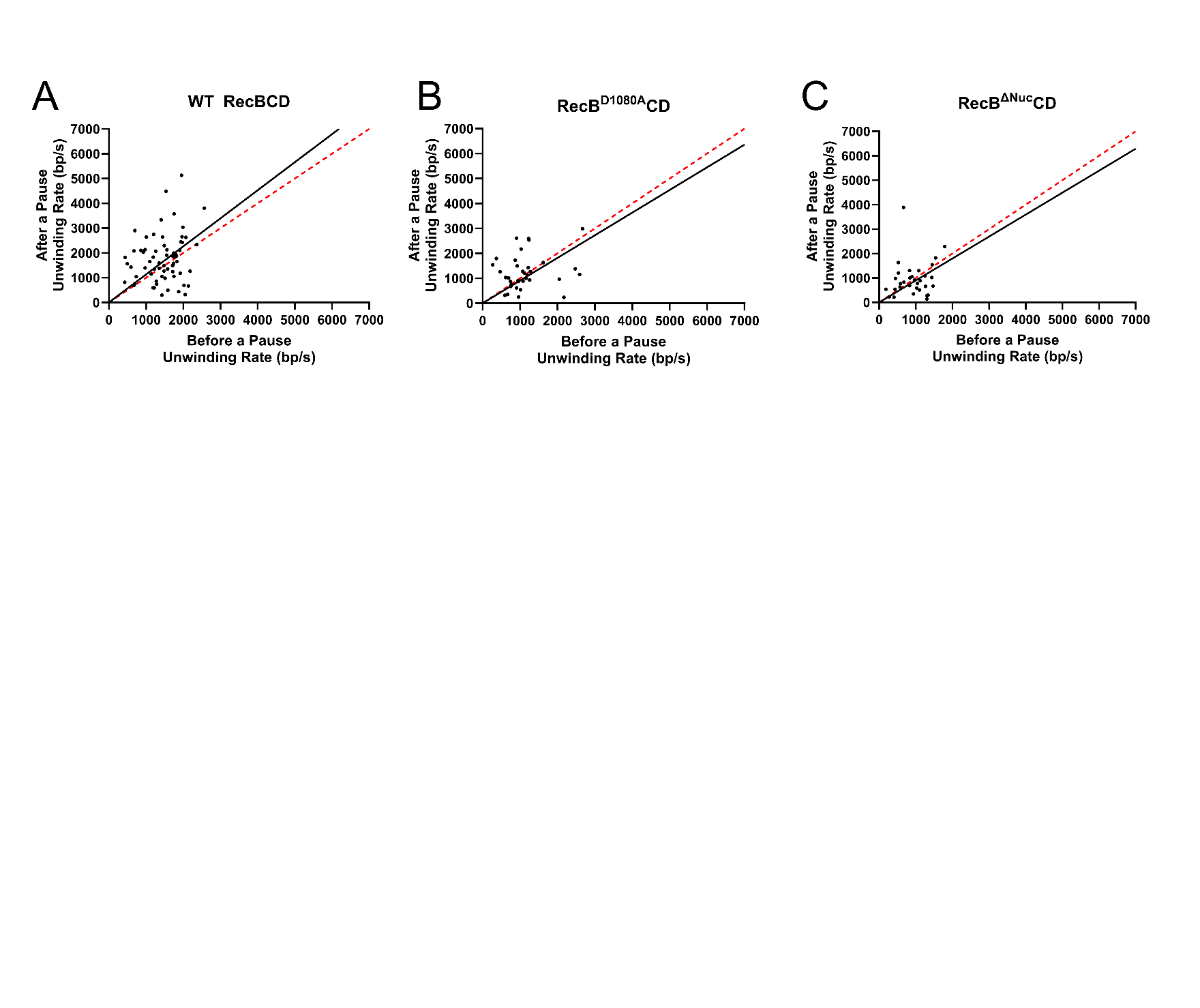

**Supplemental Figure S8. DNA unwinding rates change randomly after a DNA unwinding pause.** Rates of unwinding for RecBCD, RecB^D1080A^CD, and RecB^∆Nuc^CD before a pause and after a pause are plotted here. The data was fit to a linear line with a y-intercept constrained to the origin (black) to compare with line constrained to the origin and with a slope = 1 (red dotted). **A)** RecBCD slope = 1.13±0.17 (mean±95% CI), n = 65. **B)** RecB^D1080A^CD slope = 0.91±0.22 (mean±95% CI), n = 35.**C)** RecB^∆Nuc^CD slope = 0.90±0.27 (mean±95% CI), n = 37.

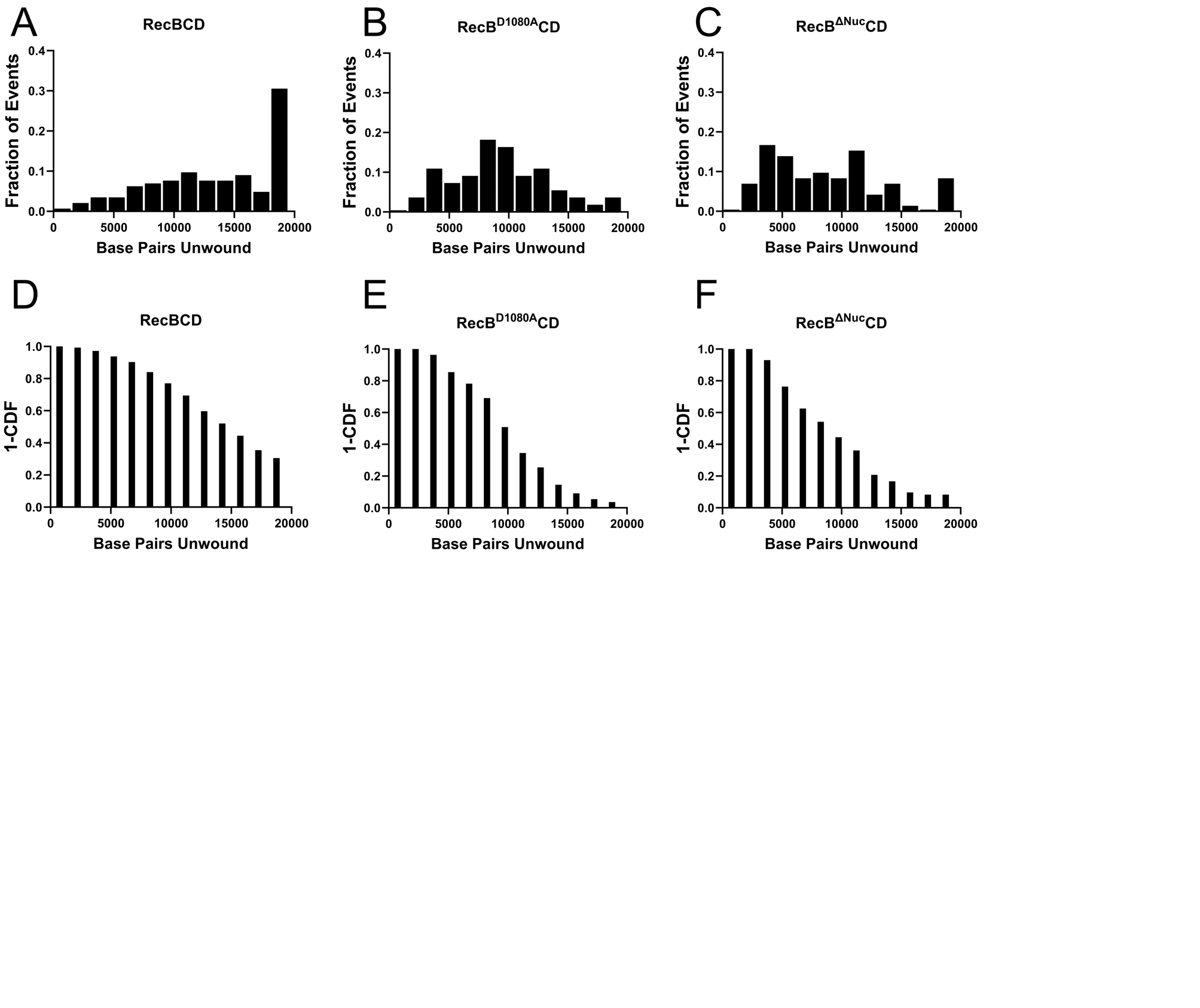

**Figure S9. DNA unwinding processivities for RecBCD, RecB^D1080A^CD, and RecB^∆Nuc^CD. A-C)** Normalized histograms of the number of bp unwound by RecBCD (A), RecB^D1080A^CD (B), and RecB^∆Nuc^CD (C). **D-F)** Characterization of the fraction of motors (1-CDF) vs. bp unwound for RecBCD (D), RecB^D1080A^CD (E), and RecB^∆Nuc^CD (F) experiments. **A-F)** RecBCD (n = 144), RecB^D1080A^CD (n = 55), and RecB^∆Nuc^CD (n = 72).

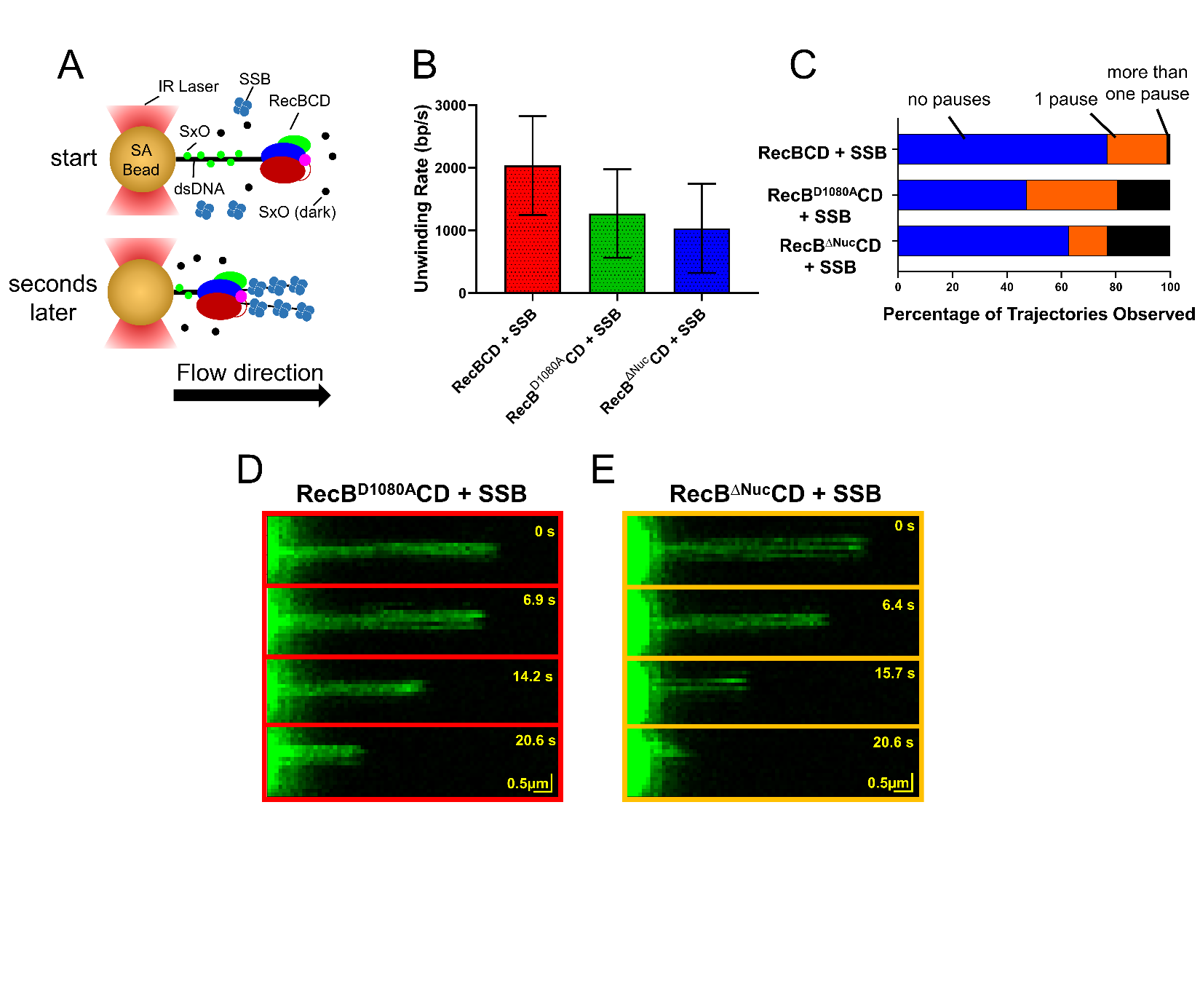

**Figure S10. *E. coli* SSB increases the rate of RecBCD dsDNA unwinding and eliminates putative ssDNA structures during RecB^D1080A^CD & RecB^∆Nuc^CD dsDNA unwinding. A)** Cartoon depicting the DNA unwinding assay in the presence of *E. coli* SSB as it coats newly formed ssDNA. **B)** Unwinding rates in the presence of 124 nM of *E. coli* SSB for WT RecBCD 2035±788 bp/s (n=106), RecB^D1080A^CD 1270±707 bp/s (n=101), and RecB^∆Nuc^CD 1033±711 bp/s (n=136) mean±SD (n). Student’s T-test reveal that both RecB^D1080A^CD & RecB^∆Nuc^CD unwinds DNA significantly slower than WT RecBCD (p-value <0.0001) and that RecB^∆Nuc^CD unwinds DNA significantly slower than RecB^D1080A^CD (p-value = 0.0075). **C)** Presented here are the fraction of RecBCD (n = 106), RecB^D1080A^CD (n =101), and RecB^∆Nuc^CD (n =136) trajectories that contain no pauses (blue), one pause (orange), and multiple pauses (black). **D & E)** Timelapse depicting the unwinding of a dsDNA tether by RecB^D1080A^CD (D) and RecB^D1080A^CD (E), without the formation of structures in the ssDNA portion of the tether in the presence of SSB.

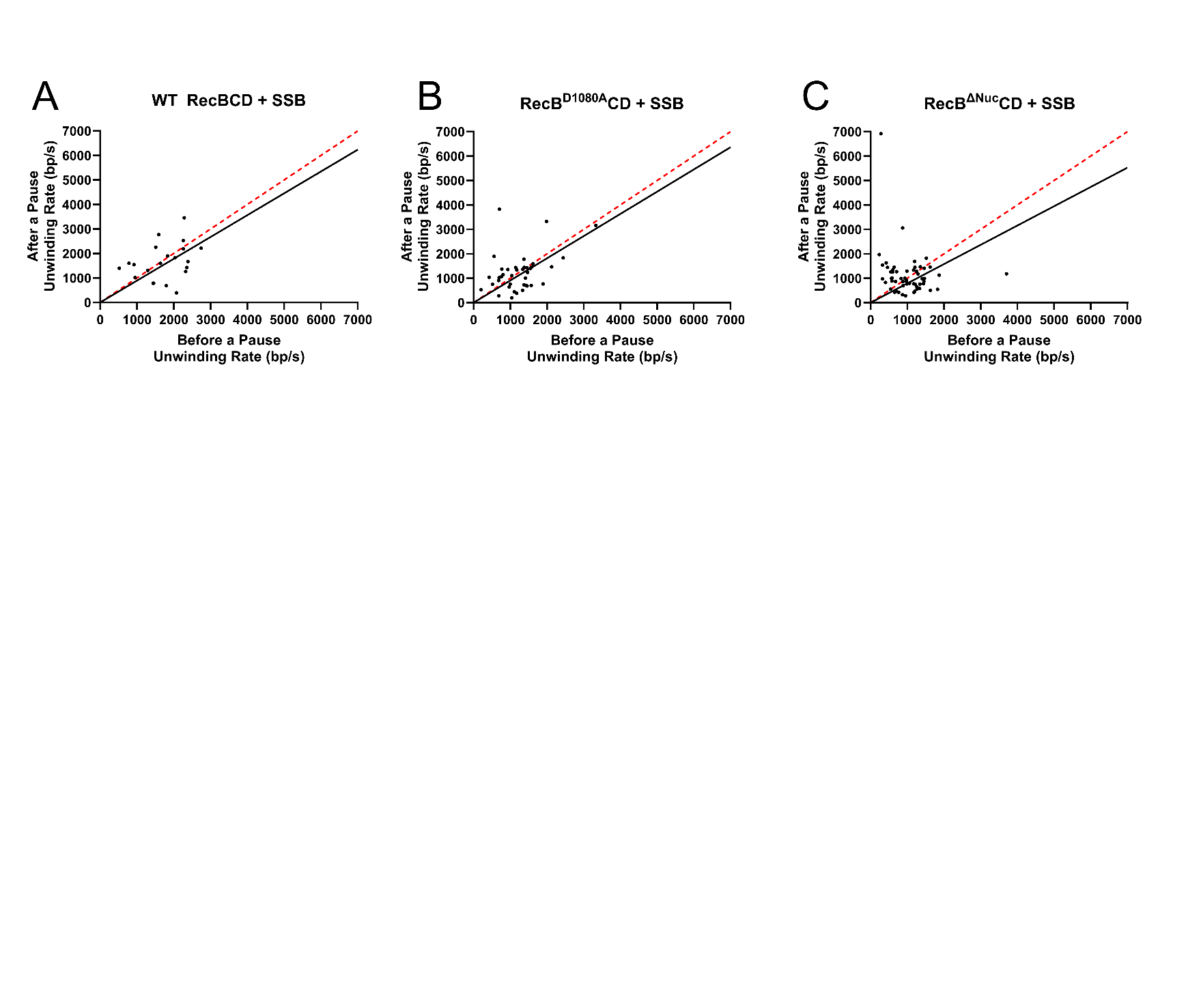

**Figure S11. DNA unwinding rates change randomly after a DNA unwinding pause in the presence of SSB protein.** Rates of unwinding for RecBCD, RecB^D1080A^CD, and RecB^∆Nuc^CD before a pause and after a pause in the presence of 124 nM *E. coli* SSB are plotted here. The data was fit to a linear line with a y-intercept constrained to the origin (black) to compare with line constrained to the origin and with a slope = 1 (red dotted line). **A)** RecBCD slope = 0.89±0.19 (mean±95% CI), n = 21. **B)** RecB^D1080A^CD slope = 0.91±0.18 (mean±95% CI), n = 39. **C)** RecB^∆Nuc^CD slope = 0.79±0.26 (mean±95% CI), n = 57.

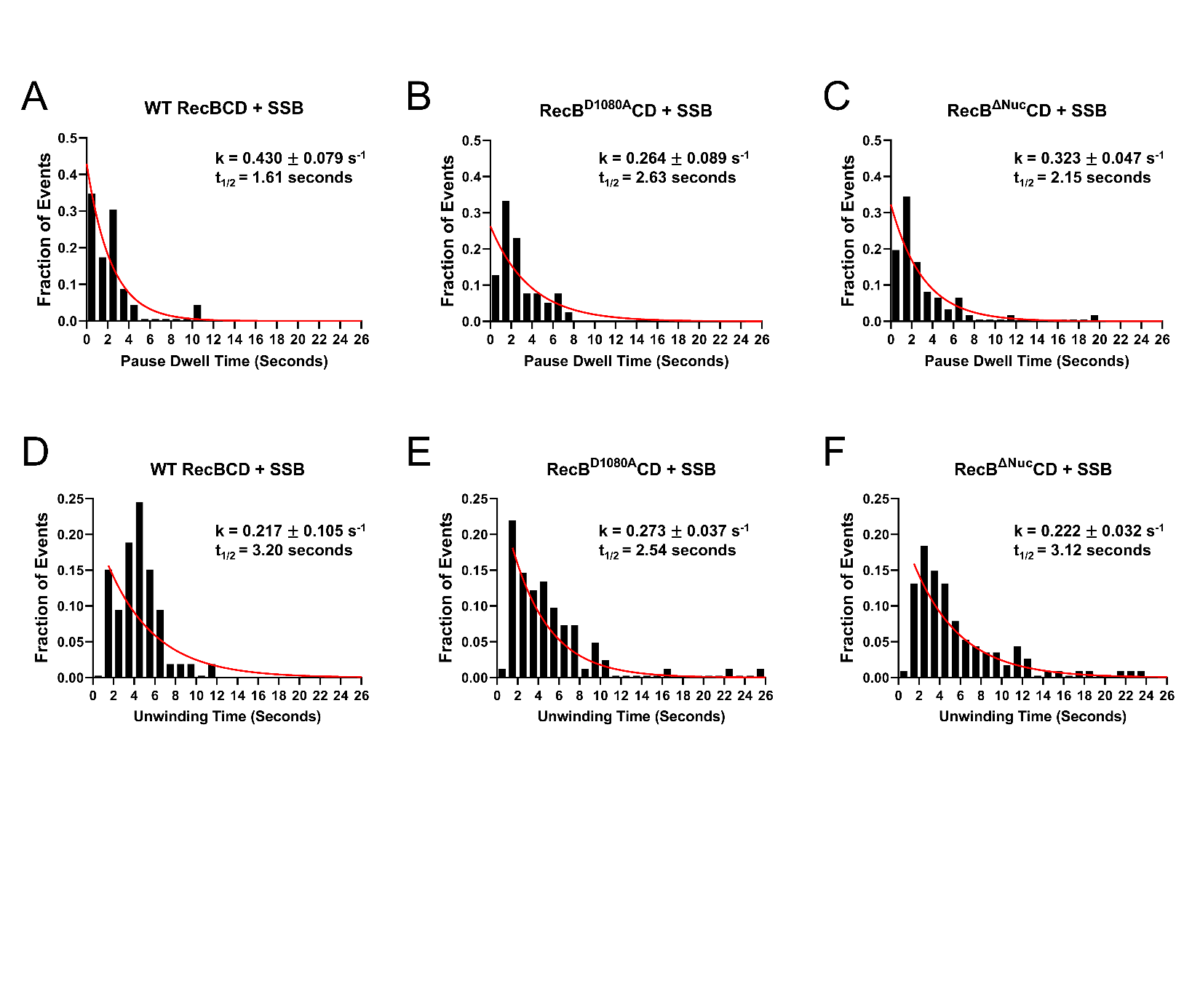

**Figure S12. Quantification of pause dwell times and DNA unwinding times in the presence of SSB protein. A-F)** Pause and unwinding dwell times for RecBCD, RecB^D1080A^CD, and RecB^∆Nuc^CD in the presence of 124 nM *E. coli* SSB were quantified and fit to an exponential probability density function in red (y=ke^-kx^). Rates (k) for each distribution are presented on the plot with the standard deviation of the fit. **A)** WT RecBCD (n = 23), **B)** RecB^D1080A^CD (n = 39), **C)** and RecB^∆Nuc^CD (n = 61), **D)** WT RecBCD (n = 53) **E)** RecB^D1080A^CD (n = 82) **F)** RecB^∆Nuc^CD (n = 114).

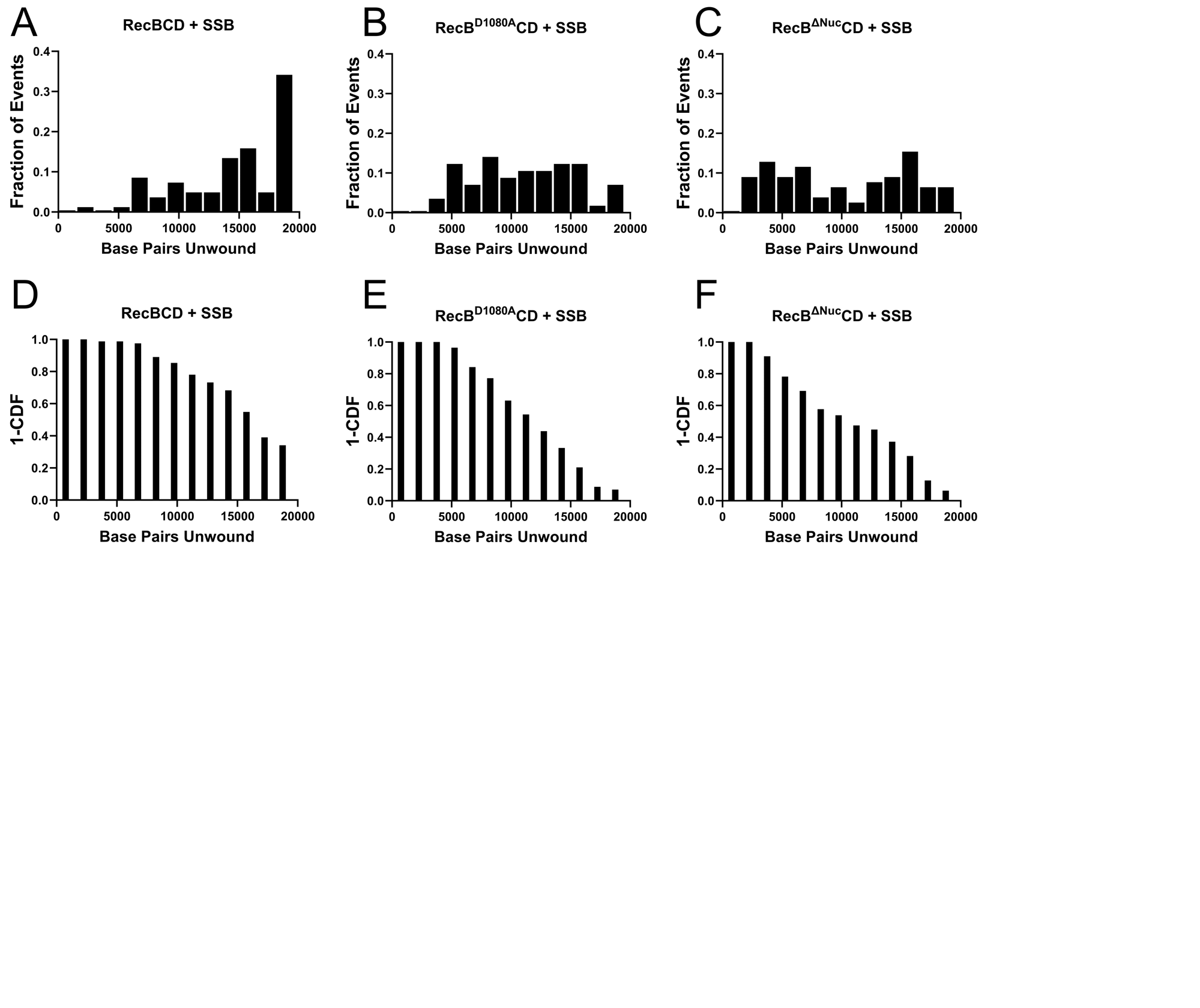

**Figure S13. Processivities of RecBCD, RecB^D1080A^CD, and RecB^∆Nuc^CD increase slightly in the presence of *E. coli* SSB. A-C)** Normalized histograms of the number of bp unwound by RecBCD + SSB (A), RecB^D1080A^CD + SSB (B), and RecB^∆Nuc^CD + SSB (C). **D-F)** Characterization of the fraction of motors engaged in unwinding (1-CDF) vs. bp unwound for RecBCD + SSB (D), RecB^D1080A^CD + SSB (E), and RecB^∆Nuc^CD + SSB (F) experiments. **A-F)** RecBCD + SSB (n = 82), RecB^D1080A^CD + SSB (n = 57), and RecB^∆Nuc^CD + SSB (n = 78).
